# Redox-active secondary metabolites act as interspecies modulators of antibiotic resilience

**DOI:** 10.1101/2021.12.01.470848

**Authors:** Lucas A. Meirelles, Dianne K. Newman

## Abstract

Bacterial opportunistic pathogens make a wide range of secondary metabolites both in the natural environment and when causing infections, yet how these molecules mediate microbial interactions and their consequences for antibiotic treatment are still poorly understood. Here, we explore the role of two redox-active secondary metabolites, pyocyanin and toxoflavin, as interspecies modulators of antibiotic resilience. We find that these molecules dramatically change susceptibility levels of diverse bacteria to clinical antibiotics. Pyocyanin is made by *Pseudomonas aeruginosa*, while toxoflavin is made by *Burkholderia gladioli*, organisms that infect cystic fibrosis and other immunocompromised patients. Both molecules alter the susceptibility profile of pathogenic species within the “*Burkholderia cepacia* complex” to different antibiotics, either antagonizing or potentiating their effects, depending on the drug’s class. Defense responses regulated by the redox-sensitive transcription factor SoxR potentiate the antagonistic effects these metabolites have against fluoroquinolones, and the presence of genes encoding SoxR and the efflux systems it regulates can be used to predict how these metabolites will affect antibiotic susceptibility of different bacteria. Finally, we demonstrate that inclusion of secondary metabolites in standard protocols used to assess antibiotic resistance can dramatically alter the results, motivating the development of new tests for more accurate clinical assessment.

## INTRODUCTION

The use of antibiotics revolutionized medicine in the 20^th^ century, but the evasion of antibiotic treatment by pathogens is a pressing health concern in the 21^st^ century (Fair and Tor, 2014; Hutchings et al., 2019; MacLean and San Millan, 2019). Without new approaches, it is estimated that by 2050 our ability to treat infections with antibiotics will dramatically decrease, resulting in up to 10 million deaths per year (O’Neill, 2016). Bacteria can withstand antibiotic treatment in many ways, including by acquiring resistance mutations and/or by developing physiological tolerance, which together enable antibiotic resilience (Blair et al., 2015; Levin-Reisman et al., 2017; Windels et al., 2019b, 2019a). Following recent terminology guidelines (Balaban et al., 2019; Brauner et al., 2016; Kester and Fortune, 2014), we define (i) ***antibiotic resistance*** as “the ability to grow in the presence of an antibiotic at a given concentration”; (ii) ***antibiotic tolerance*** as “the ability to survive transient antibiotic exposure”; and (iii) ***antibiotic resilience*** as “the ability of a bacterial population to be refractory to antibiotic treatment” (Perry et al., 2021). When physicians face situations where treatments do not work in the clinic, this is typically caused due to increased tolerance and/or resistance levels (Windels et al., 2019b).

One important and underappreciated factor that promotes both resistance and tolerance is the production of secondary metabolites (Perry et al., 2021). These are a broad category of molecules generally produced under slow-growth conditions (i.e., “stationary phase” in the laboratory batch cultures) that are secreted extracellularly or kept inside the cell (Davies, 2013; Price-Whelan et al., 2006). When secreted, secondary metabolites have the potential to affect their surroundings, including other microbes (Davies, 2013; Tyc et al., 2017). Even though it is well known that microbes can influence each other through secondary metabolite production (Tyc et al., 2017), only recently has this concept been considered in the context of antibiotic susceptibility (Meirelles et al., 2021; Perry et al., 2021; Radlinski et al., 2017). Secondary metabolites typically affect antibiotic susceptibility by (i) inducing efflux systems or (ii) modulating redox homeostasis or oxidative stress responses (Meirelles et al., 2021; Perry et al., 2021). It stands to reason that they might significantly affect our ability to treat polymicrobial infections.

Certain secondary metabolites are redox-active molecules that promote multifaceted benefits for their producers, from acquiring nutrients to controlling redox homeostasis and promoting anaerobic survival (Glasser et al., 2014; Jo et al., 2017; Price-Whelan et al., 2006). Because they readily react with oxygen (leading to reactive oxygen species generation) and with Fe-S clusters inside the cells (directly oxidizing proteins), redox-active secondary metabolites also display high toxicity levels (Gu and Imlay, 2011; Imlay, 2013; Laursen and Nielsen, 2004; Singh et al., 2013). For this reason, they are often called “natural antibiotics”. Their toxicity puts selective pressure on producers and other microbes that are commonly found together with producing species to develop defense mechanisms against their toxic effects (Davies and Davies, 2010; Martinez, 2009; Waglechner et al., 2019). We have recently proposed that, through the induction of such defense mechanisms, redox-active metabolites can promote collateral resilience to certain clinical drugs (Meirelles et al., 2021). This is particularly relevant when the redox-active metabolite shares structural similarities to the drug used (Meirelles et al., 2021). It is still unclear, however, how broad this phenomenon is when considering distinct redox-active secondary metabolites, the molecular mechanisms involved, and the drugs affected.

Here, we explore the role of two redox-active secondary metabolites as modulators of antibiotic resilience. We focus on pyocyanin and toxoflavin, structurally similar compounds made by diverse opportunistic pathogens, including *Pseudomonas aeruginosa* and *Burkholderia* species. Our mechanistic investigation of representatives of strains that are commonly co-isolated from clinical infections reveals that the production of these metabolites and conserved machinery that senses them can affect the antibiotic susceptibility levels of neighboring species in a significant and predictable fashion. Our findings motivate the development of new approaches to improve the accuracy of antibiotic resistance diagnostics, which is necessary to optimize the use of available drugs.

## RESULTS

### Pyocyanin produced by *Pseudomonas aeruginosa* induces complex defense responses in *Burkholderia*

To investigate how redox-active secondary metabolite producers might affect antibiotic susceptivity levels in polymicrobial infections, we started by exploring interactions between two opportunistic pathogens relevant to the cystic fibrosis (CF) lung environment: *Pseudomonas aeruginosa* and *Burkholderia multivorans*. *P. aeruginosa* is a global opportunist pathogen that causes serious infections in patients with CF, chronic wounds, and compromised immune systems (Driscoll et al., 2007). *B. multivorans* is part of the *Burkholderia cepacia* complex (Bcc). Species in this group can cause severe chronic infections and are associated with dire prognoses in patients with CF (Coutinho et al., 2011; Lipuma, 2010). Even though *P. aeruginosa* and Bcc species already display high levels of antibiotic resistance (Driscoll et al., 2007; Rhodes and Schweizer, 2016), recent evidence indicated that the production of pyocyanin (PYO) by *P. aeruginosa* can modulate susceptibility levels to fluoroquinolone antibiotics in Bcc (Meirelles et al., 2021) (Fig. 1A), but the mechanisms involved in this process were unknown. PYO is part of a diverse group of molecules classified as phenazines, which are important virulence factors during *P. aeruginosa* infections and have been detected in patients (Cruickshank and Lowbury, 1953; Lau et al., 2004; Wilson et al., 1988).

**Figure 1.**
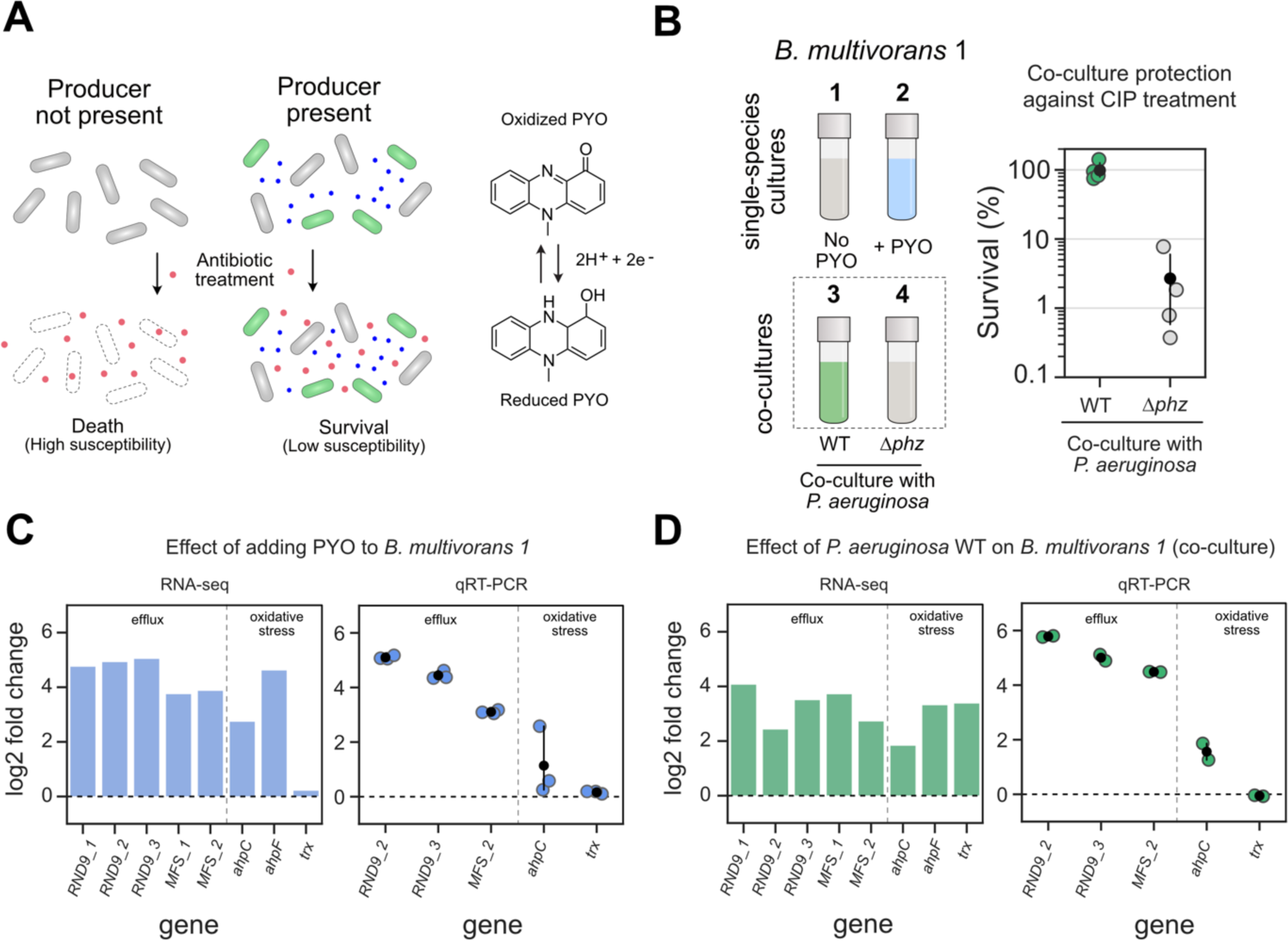
PYO produced by *P. aeruginosa* induces complex defense responses in *B. multivorans.* **A.** Left: Model of secondary metabolite-mediated induction of survival against antibiotics in microbial populations. Blue dots represent secondary metabolites made by producer species (green cells); red dots represent the antibiotic. Right: one example of such metabolites is PYO, a redox-active molecule produced by *P. aeruginosa*. **B.** Left: Conditions used during RNA-seq and qRT-PCR experiments (1 to 4). In all cases, responses were measured using *B. multivorans* 1 WT (see Materials and Methods). The strain was either grown as a single-species culture and exposed or not to PYO (conditions 1 and 2), or co-cultured with *P. aeruginosa* that can (WT) or cannot (Δ*phz*) make the phenazine (conditions 3 and 4). Right: *B. multivorans* 1 tolerance against ciprofloxacin (CIP, 10 µg/mL) when in co-culture with WT or Δ*phz P. aeruginosa* (n = 4). **C-D.** Representative genes highlighting the responses induced by exogenously added PYO (C) or by co-culturing with PYO-producing *P. aeruginosa* (D). RNA-seq and qRT-PCR results are shown as bar (left) and strip (right) plots, respectively. Genes are displayed in two categories (efflux and oxidative stress) and named accordingly to their draft annotation or to their respective homolog in the *P. aeruginosa* genome. For qRT-PCR data, n = 3 in panel C, and n = 2 in panel D (see “experiment 1” in the Material and Methods). For full dataset for each comparison that includes transcriptional changes across *B. multivorans* genome and their respective loci tags, see Table S2. For additional qRT-PCR results, see Fig. S1.

To screen PYO-mediated molecular responses induced in *B. multivorans*, we first performed RNA-seq experiments either by (i) exposing the cells to exogenously added PYO or (ii) by co-culturing *B. multivorans* with *P. aeruginosa* (Fig. 1B-left). Control treatments involved sequencing RNA of *B. multivorans* alone or in co-culture with a *P. aeruginosa* strain that cannot make phenazines (Δ*phz*), including PYO. We used the *B. multivorans* strain AU42096 (referred to here as *B. multivorans* 1, see Table S1 for details). Notably, when in the presence PYO-producing *P. aeruginosa*, *B. multivorans* 1 is highly tolerant of fluoroquinolones such as ciprofloxacin (Fig. 1B-right). We found that PYO, either when added directly or when produced by *P. aeruginosa*, induces a set of responses in *B. multivorans* 1 (Table S2) that can be grouped into two broad classes: (i) induction of specific efflux systems, including a resistance-nodulation-division (RND) efflux system and a potential major facility superfamily (MFS) transporter, and (ii) oxidative stress responses, including alkyl hydroperoxide reductases (Fig. 1C). These results were confirmed by qRT-PCR (Fig. 1C-D). Not surprisingly, exposing *B. multivorans* to *P. aeruginosa* lead to more complex transcriptional responses than exposure to PYO alone, likely due to the wide variety of molecules secreted by *P. aeruginosa* (Bartell et al., 2017).

Despite the complexity of the *B. multivorans* transcriptomic responses, the induction of one operon called our attention: an efflux system commonly known as RND-9 (Podnecky et al., 2015). All the genes of this operon (Bmul_3930, Bmul_3931, and Bmul_3932) were induced in the presence of PYO, either when added exogenously or via production by co-cultured *P. aeruginosa* WT (Figs. 1C-D, Fig. S1, Table S2). Given the importance of RND efflux systems in antibiotic tolerance and resistance (Li et al., 2015), especially when these systems are induced by PYO (Meirelles et al., 2021), we were motivated to investigate whether the RND-9 efflux system in *B. multivorans* and other *Burkholderia* species is required to confer antibiotic tolerance in the presence of PYO.

### Redox-regulated efflux mediates *Burkholderia* susceptibility to pyocyanin and its collateral effects on antibiotic resilience

The PYO-mediated induction of RND-9 in *Burkholderia multivorans* seen in our RNA-seq and qRT-PCR experiments led us to investigate this efflux system more carefully. When searching the genomic region where RND-9 is present, we noticed a flanking gene annotated as a “MerR family transcriptional regulator” (Bmul_3929, Fig. 2A). BLASTing the Bmul_3929 protein sequence from our *B. multivorans* 1 strain against proteins encoded in the *P. aeruginosa* PA14 and *P. aeruginosa* PAO1 genomes (Winsor et al., 2016) resulted in their respective SoxR as first hits (gene locus tags PA2273 and PA14_35170, respectively, Table S3). SoxR is primarily studied in *P. aeruginosa* and *E. coli*; in both cases, the transcription factor contains a Fe-S cluster that can be directly oxidized by redox-active molecules, such as PYO (Fig. 2A) (Dietrich et al., 2006; Gu and Imlay, 2011). This results in an induction of stress responses, including efflux systems (Dietrich et al., 2006; Meirelles et al., 2021; Meirelles and Newman, 2018). However, the role of SoxR in other organisms is much less understood.

**Figure 2.**
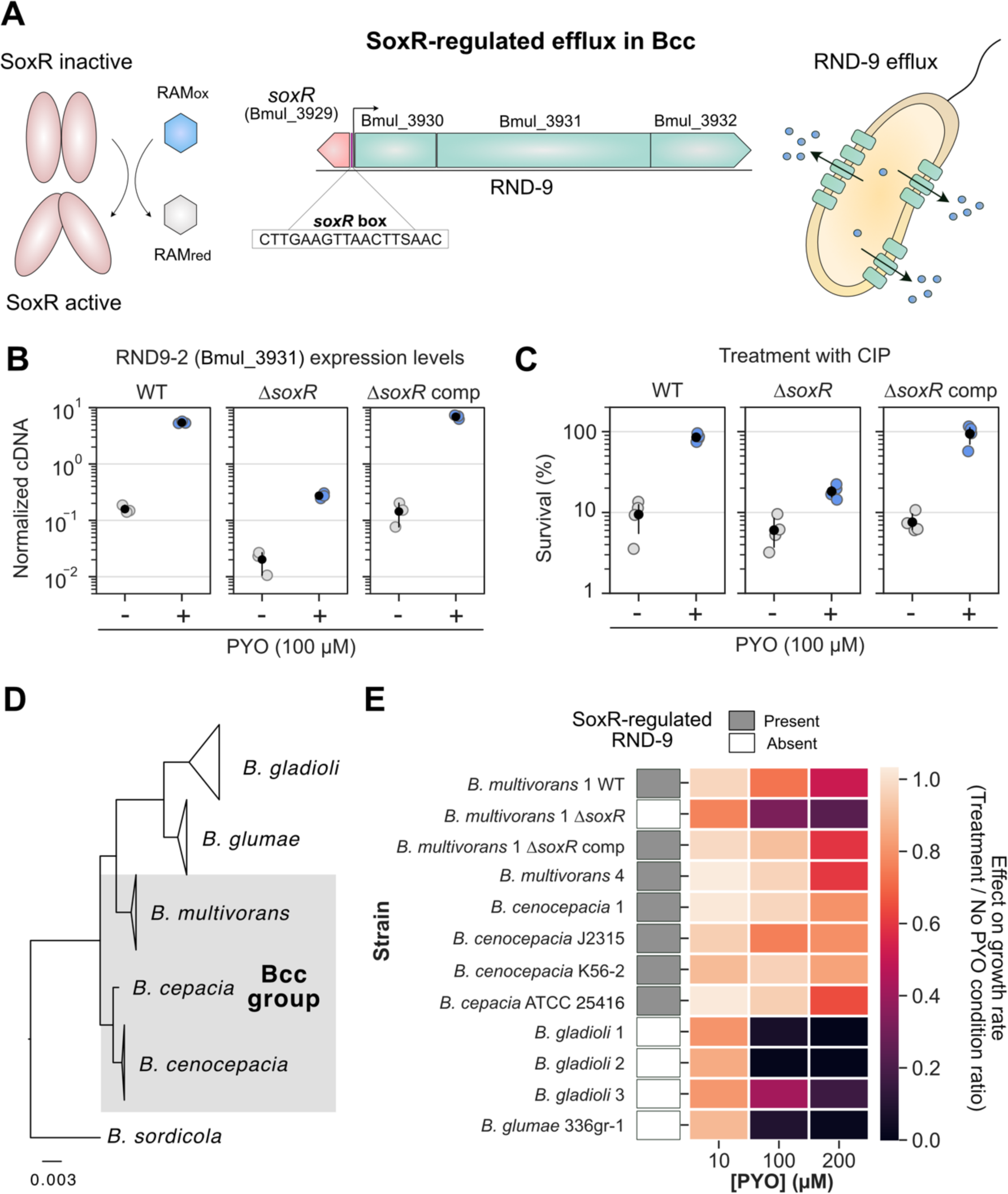
Redox-regulated efflux mediates *Burkholderia* susceptibility to PYO and its collateral effects on antibiotic resilience. **A.** SoxR-mediated regulation of RND-9 in Bcc. The example is based on the *B. multivorans* ATCC 17616 genome locus structure and orientation. SoxR is oxidized by redox active metabolites (RAMs), triggering its activation (left) (Dietrich et al., 2008; Imlay, 2013; Perry et al., 2021). In Bcc, SoxR is commonly found adjacent to the RND-9 efflux system; the SoxR box (Dietrich et al., 2008) is found upstream to RND-9 genes, and the sequence displayed is the consensus found in this genomic region for the Bcc strains studied (center). Following other examples of RND efflux systems, proteins derived from Bmul_3930 and Bmul_3931 are likely associated with the inner membrane, while the one from Bmul_3932 is likely an outer membrane protein (Du et al., 2018). SoxR-mediated induction of the system allows efflux of toxic molecules by the bacterial cell (right). **B.** Expression levels of the second gene in the RND-9 operon (Bmul_3931) measured by qRT-PCR in different *B. multivorans* strains in the presence or absence of PYO (n = 3). Δ*soxR* comp means complementation of Δ*soxR*. Data is shown as normalized cDNA (see Materials and Methods). For additional qRT-PCR results, see Fig S2. **C**. Effect of PYO on tolerance to ciprofloxacin (CIP, 10 μg/mL) in the same three *B. multivorans* strains (n = 4). **D**. Phylogenetic relationship between the *Burkholderia* species used in this study (gray shading highlights species within the Bcc group). For full tree detailing different strains, see Fig. S4. For broader phylogenetic placement of these species within the *Burkholderia* genus, see Depoorter et al. (2016). **E.** PYO effect on growth rates of distinct strains of the different *Burkholderia* species studied. Data for three different concentrations are shown (10, 100 and 200 µM). The results are shown as a ratio of the growth rates for each strain under different PYO concentrations by their growth rates in the “No PYO” condition (i.e. values close to 1 mean no inhibition, while values close to 0 mean severe growth inhibition by PYO). Presence or absence of the genomic locus containing SoxR and RND-9 in these strains is indicated by the grey or white boxes, respectively. Growth rates were estimated based on growth curves under the different conditions (Fig. S5). In panels B and C, the black dots mark the means and error bars represent 95% confidence intervals.

First, we hypothesized that, by sensing redox-active metabolites such as PYO, SoxR might regulate the expression of RND-9, which is then used to export them. We confirmed that in our model Bcc strain, *B. multivorans* 1, the RND-9 promoter region has a SoxR box (Fig. 2A). The SoxR box has been shown to be a strong predictor for the operon’s regulation by SoxR (Dietrich et al., 2008), suggesting that SoxR regulates RND-9 expression in *B. multivorans*. To confirm that SoxR is necessary for PYO-mediated increase in RND-9 expression in *B. multivorans* 1, we made a Δ*soxR* deletion mutant. As predicted, PYO did not increase RND-9 expression in the Δ*soxR* strain to the extent it did in the WT, and overall expression levels were significantly lower and comparable to background levels (i.e., no PYO in the WT) (Fig. 2B, Fig. S2). Complementation of the *soxR* gene restored PYO-mediated activation of RND-9 (Fig. 2B, Fig. S2).

We hypothesized that the PYO-mediated increase in resilience against fluoroquinolones in *B. multivorans* was mediated by SoxR-induction of RND-9. Supporting our hypothesis, the Δ*soxR* strain was much less tolerant to ciprofloxacin in the presence of PYO than the WT (Fig. 2C). Even though there was an apparent slight increase in the survival percentage for Δ*soxR* when PYO is present, we attribute this to a normalization artifact: because PYO affected the growth of Δ*soxR*, the strain reached a lower cell density when PYO was present, causing the normalization (i.e., % survival) to be calculated by a smaller CFU number. Raw CFU values showed that PYO only mildly increased the number of surviving cells upon treatment with ciprofloxacin in the Δ*soxR* strain, while the effect was significantly higher in the WT strain (Fig. S3). Complementation of *soxR* in the mutant restored WT levels of tolerance (Fig. 2C and Fig. S3).

Because PYO significantly increased the expression of RND-9 in *B. multivorans*, we decided to investigate the importance of this system for PYO resistance in other *Burkholderia* species. Specifically, we hypothesized that the presence of the SoxR-regulated RND-9 efflux system might be used to predict how well different *Burkholderia* species can manage PYO toxicity. Export is crucial for handling PYO’s toxic effects (Meirelles et al., 2021; Meirelles and Newman, 2018). In this scenario, *Burkholderia* species containing SoxR-mediated RND-9 would sense the presence of PYO secreted by *P. aeruginosa* and quickly induce the efflux system, increasing their fitness in the presence of the molecule. We searched the *Burkholderia* Genome Database (www.burkholderia.com) (Winsor et al., 2008) for species commonly found in CF patients as well as in the environment (e.g., plant-associated strains). These included species within the Bcc (such as *B. multivorans*, *B. cepacia*, *B. cenocepacia*) and non-Bcc species (such as *B. glumae* and *B. gladioli*). We obtained several strains within these species, many of which are clinical isolates from lung respiratory infections (see Table S1), compared their phylogenetic relationship (Fig. 2D, Fig. S4), and assessed whether the SoxR-regulated RND-9 efflux system was present (Fig. 2E). The system’s presence was determined by (i) directly inspecting the genome (available in databases or through whole-genome sequencing), by (ii) PCR-amplifying a fragment containing part of the SoxR/RND-9 genome locus, or by (iii) by inference based on the genomic content of closely-related strains within the same species in genome databases, such as the *Burkholderia* Genome Database (Winsor et al., 2008) and Integrated Microbial Genomes and Microbiomes IMG/M (Chen et al., 2021; Mukherjee et al., 2021) (see Materials and Methods for details). In parallel, we measured the growth rates of these strains in the presence of different concentrations of PYO (Fig. 2E and Fig. S5). We saw a strong correlation between the presence of SoxR/RND-9 locus and the strain’s ability to handle PYO toxicity. The system was present in all Bcc species tested, and growth of these strains was only mildly affected by PYO, even at concentrations as high as 200 µM (Fig. 2E and Fig. S5). However, growth of strains lacking the regulator/efflux system (*B. glumae*, different *B. gladioli* strains, and the *B. multivorans* Δ*soxR*) was dramatically inhibited by PYO (Fig. 2E and Fig. S5). This indicates that SoxR-regulated efflux plays a conserved role in how well *Burkholderia* species can tolerate PYO, potentially affecting their ability to inhibit habitats where *P. aeruginosa* is present. Moreover, this system also modulates *P. aeruginosa*’s impact on *Burkholderia* susceptibility to antibiotics because the *P. aeruginosa*-mediated increase in antibiotic tolerance on *B. multivorans* is SoxR dependent (Fig. S6).

Overall, these results highlight the importance of SoxR and its redox-regulated efflux systems during interspecies interactions mediated by redox-active secondary metabolites, demonstrating their antagonistic effect on fluoroquinolone susceptibility.

### Redox-active secondary metabolites as interspecies modulators of antibiotic susceptibility: the toxoflavin example

Our results with PYO motivated us to ask whether other redox-active secondary metabolites might similarly be capable of modulating antibiotic susceptibility in polymicrobial infections (Perry et al., 2021). Accordingly, we searched for other potential candidate molecules known to be made by pathogens that have been found infecting CF patients. We found that phenazine-1-carboxylic acid (PCA), another phenazine made by *P. aeruginosa*, also dramatically increased *B. multivorans* tolerance against ciprofloxacin in a SoxR-dependent manner (Fig. S7). But to test the generality of the phenomenon, we looked for non-phenazine molecules made by species other than *Pseudomonas*. One example we found was toxoflavin (TOX) (Fig. 3A). TOX is made by different *Burkholderia* species, including *B. glumae* and *B. gladioli*, and has been proposed to increase the fitness and virulence of its producers when infecting plants (Chen et al., 2012; Jeong et al., 2003; Lee et al., 2016). *B. glumae* is a plant pathogen, while *B. gladioli* can cause disease in plants and humans (Ham et al., 2011; Jones et al., 2021; Lipuma, 2010). In fact, *B.gladioli* is among the most prevalent *Burkholderia* species in CF patients (Lipuma, 2010). Though TOX detection in CF sputum has not been attempted to our knowledge, clinical *B. gladioli* strains commonly produce TOX *in vitro* (Jones et al., 2021). TOX is redox-active and thought to induce efflux and oxidative stress response in bacteria (Fig. 3A-B) (Latuasan and Berends, 1961; Stern, 1935). Therefore, TOX is a potentially clinically-relevant molecule and a good candidate for testing the hypothesis that redox-active secondary metabolites have broad potential to modulate antibiotic resilience.

**Figure 3.**
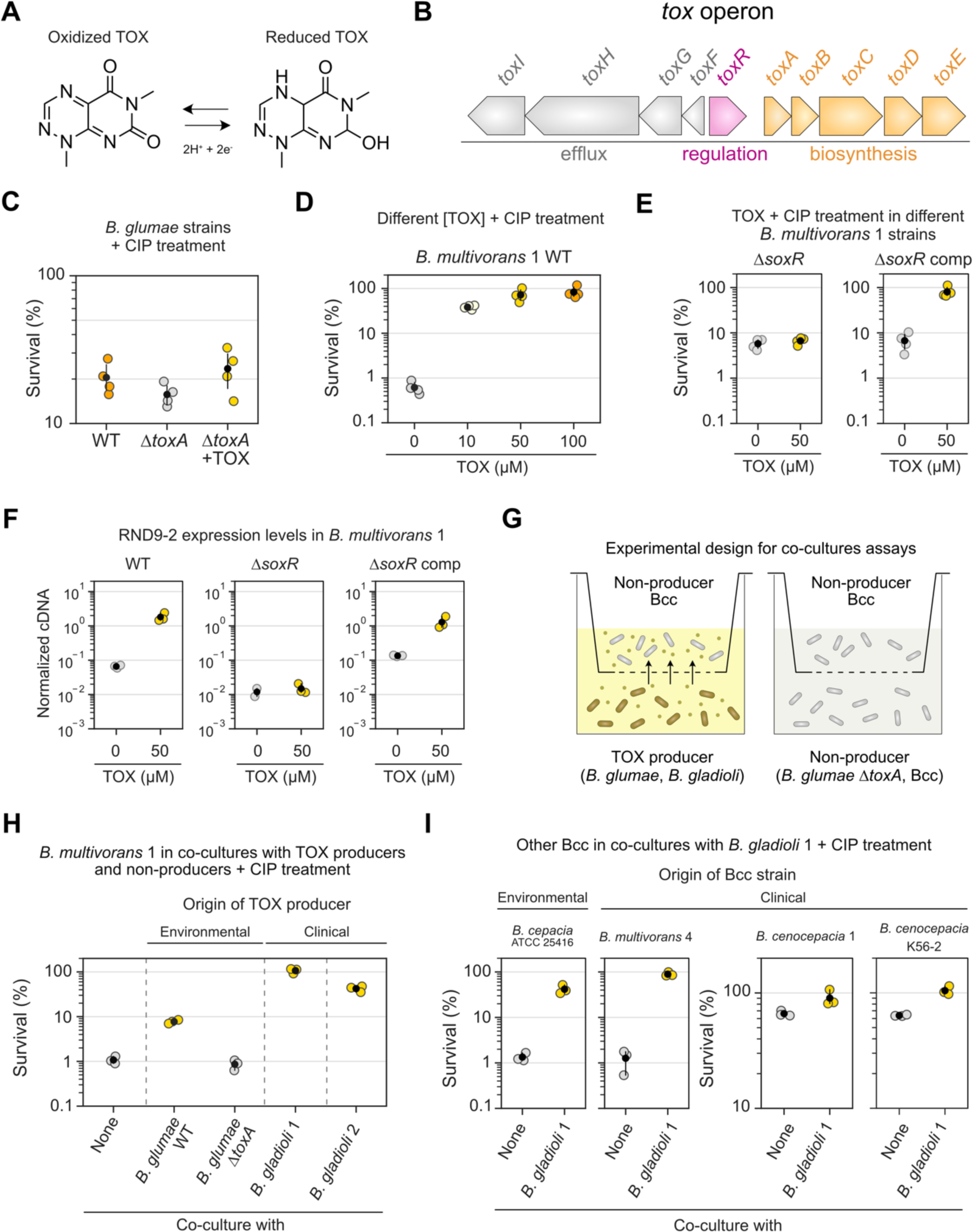
TOX increases tolerance against ciprofloxacin in Bcc. **A.** TOX molecule and its redox states. **B.** Genomic structure of the *tox* operon present in the TOX producer *B. glumae* (Chen et al., 2012; Kim et al., 2004). Similar genomic context is found in other TOX producers (Jones et al., 2021). **C.** Effect of TOX on tolerance to ciprofloxacin (CIP, 1 µg/mL) in different *B. glumae* strains (n = 4). TOX was either produced endogenously by the WT strain or added exogenously (50 µM) to the Δ*toxA* mutant. **D.** Effect of different TOX concentrations on tolerance to CIP (10 µg/mL) in the *B. multivorans* 1 WT strain (n = 4). **E.** Effect of TOX on tolerance to CIP (10 µg/mL) in the *B. multivorans* 1 Δ*soxR* or Δ*soxR* comp strains (n = 4). This experiment was performed separately from that shown in panel D. **F.** Expression levels of the second gene in the RND-9 operon (Bmul_3931) measured by qRT-PCR in different *B. multivorans* strains in the presence or absence of TOX (n = 3 for all except “Δ*soxR* + 0 µM TOX”, where n = 2). Data is shown as normalized cDNA (see Materials and Methods). Also, see Figs. S9 additional qRT-PCR data. **G.** Experimental design used during the co-culture antibiotic assays (see Materials and Methods for details). **H.** Effect of TOX produced by different producer species (*B. glumae* WT, and several *B. gladioli* strains) on the tolerance of *B. multivorans* 1 to CIP (10 µg/mL). Conditions where TOX was not produced were: “none” (i.e. only *B. multivorans* 1 is present) or co-culture with the *B. glumae* Δ*toxA* strain that cannot make TOX. *B. glumae* was originally isolated from environmental sample (Chen et al., 2012), while *B. gladioli* strains are derived from CF patients (see Table S1). Only the top part of the co-culture plate (i.e. containing *B. multivorans* 1) was plated for CFUs for survival assessment (n = 3). **I.** Effect of TOX produced by *B. gladioli* 1 on the tolerance of multiple Bcc species (isolated from environmental and clinical samples) to CIP (10 µg/mL, n = 3). Note that the scales on the two plots assessing tolerance in *B. cenocepacia* (two on the right) are different from the ones on the left, since the background tolerance levels in these strains are much higher than the ones in *B. cepacia* or *B. multivorans* 4 (left). In panels C-F and H-I, the black dots mark the means and error bars represent 95% confidence intervals. Panel G was adapted from Meirelles et al. (2021), CC BY 4.0 (https://creativecommons.org/licenses/by/4.0/).

We started by determining whether TOX can increase ciprofloxacin tolerance in the producing species *B. glumae,* just like PYO does in *P. aeruginosa* (Meirelles et al., 2021; Schiessl et al., 2019). TOX induces the efflux system ToxFGHI in *B. glumae* (Kim et al., 2004), and we confirmed that the WT strain makes TOX under the studied conditions (Fig. S8). However, when comparing WT ciprofloxacin survival levels to a Δ*toxA* strain that cannot make TOX (Lelis et al., 2019), we did not see a significant increase in tolerance (Figs. 3C); similarly, adding TOX exogenously to Δ*toxA* did not significantly increase tolerance (Fig. 3C). Although we observed a slight tolerance increase trend when TOX was present, the magnitude of the effect is much smaller than what PYO provides to *P. aeruginosa* (Meirelles et al., 2021), revealing that ToxFGHI and the overall responses induced by TOX in *B. glumae* do not confer ciprofloxacin tolerance to this strain. Even though TOX does not provide significant protection against ciprofloxacin in *B. glumae*, we reasoned that it might act as an interspecies modulator of antibiotic resilience in another organism. For example, *B. gladioli* strains (potentially TOX producers) (Graves et al., 1997; Jones et al., 2021) can infect CF patients together with other Bcc pathogens (Kennedy et al., 2007). We thus decided to test if Bcc species might benefit from the presence of TOX when treated with ciprofloxacin. Tolerance assays using our Bcc model organism, *B. multivorans* 1, showed that TOX increased its survival against ciprofloxacin in a concentration-dependent manner (Fig. 3D). Levels of 50 µM or above made the strain completely tolerant to the antibiotic treatment (Fig. 3D). Importantly, concentrations of TOX within the 10-100 µM range are physiologically relevant since they fall within the amounts produced by *B. gladioli* and *B. glumae* under the studied conditions (Fig. S8). We next hypothesized that this tolerance phenotype is mediated by the SoxR-regulated RND-9 efflux system, like it is for PYO. Consistent with this prediction, the TOX-mediated increase in tolerance against ciprofloxacin disappeared in the Δ*soxR* strain but was restored in the SoxR-complemented strain (Fig. 3E). Moreover, using qRT-PCR to quantify the expression of RND-9 in *B. multivorans* 1, we observed that TOX increased transcription of RND-9 in *B. multivorans* 1 and its induction disappeared in the Δ*soxR* mutant but was restored in the SoxR-complemented strain (Fig. 3F, Fig. S9). This indicates that SoxR in *B. multivorans* 1 works as a broader sensor for a wide range of redox-active molecules, with direct consequences for antibiotic efficacy.

We next evaluated the effect of TOX in co-culture assays (Fig. 3G). First, we tested if *B. multivorans* 1 becomes more tolerant to ciprofloxacin when grown with environmental or clinical TOX producers (Fig. 3H). Co-culture with *B. glumae* WT (i.e., TOX present) increased *B. multivorans* 1 tolerance to ciprofloxacin. The phenotype disappeared when the strain was co-cultured with the *B. glumae* Δ*toxA* (i.e., no TOX). Notably, co-culture with two different clinical strains of *B. gladioli* isolated from CF patients dramatically increased tolerance against ciprofloxacin by *B. multivorans* 1 (Fig. 3H). These two clinical strains produce TOX (Fig. S8), and the molecule was present in the co-cultures, evident by its yellow pigmentation. To evaluate the generality of these results, we compared the ciprofloxacin tolerance of multiple Bcc strains from different species (*B. cepacia*, *B. multivorans*, and *B. cenocepacia*) to that seen when they were co-cultured with a TOX-producing clinical strain of *B. gladioli*. These Bcc strains were derived either from environmental samples or from patients. In all cases, co-culture with *B. gladioli* increased ciprofloxacin tolerance levels in the Bcc species (Fig. 3I). All co-cultures were yellow, indicating TOX was present. The effect was dramatic for *B. cepacia* and *B. multivorans*, increasing tolerance levels more than an order of magnitude; although *B. cenocepacia* strains showed higher background tolerance levels when grown alone in the absence of TOX, co-culturing these strains with TOX-producing *B. gladioli* still made them more tolerant of ciprofloxacin (Fig. 3I).

Overall, our results show that phenazines made by *P. aeruginosa* are not the only redox-active secondary metabolites that can modulate antibiotic resilience. TOX, a redox-active molecule produced by different *Burkholderia* species, can do the same, suggesting generality for the phenomenon.

### Assessing the effects of redox-active secondary metabolites on antimicrobial susceptibility testing

These mechanistically-oriented laboratory results raised an important practical question: how might redox-active secondary metabolites impact standard clinical antibiotic susceptibility testing (AST)? Current AST methods are blind to potential modulating effects of redox-active secondary metabolites. This happens because the synthesis of these metabolites is usually controlled by quorum sensing, being made only at cells densities that are much higher than what is used for typical AST inocula (Davies, 2013; Jorgensen and Ferraro, 2009). However, redox-active secondary metabolites have been detected in infections (Cruickshank and Lowbury, 1953; Wilson et al., 1988) and presumably could dramatically change the performance of certain drugs during treatment (Perry et al., 2021). To take this into account, we modified the protocol of a traditional AST assay that determines the minimum inhibitory concentration (MIC) for specific drugs. Using PYO and TOX as examples, we tested how these metabolites affect susceptibility to several antibiotics from different classes.

Our experimental design is shown in Fig. 4A and Fig. S10A. In brief, MIC assays following the EUCAST guidelines (see Materials and Methods) were performed in the absence or presence of exogenously added PYO or TOX. For these assays, we used our model organism, *B. multivorans* We tested several different (sub)classes of drugs, including fluoroquinolones (ciprofloxacin and levofloxacin), tetracyclines (tetracycline and doxycycline), amphenicols (chloramphenicol), sulfonamides (sulfamethoxazole in combination with trimethoprim), aminoglycosides (tobramycin), carbapenems (meropenem), cephalosporins (ceftazidime), and polymyxins (colistin). PYO and TOX altered the MIC of *B. multivorans* for several types of drugs. Both PYO and TOX had overall antagonistic effects on fluoroquinolones, tetracyclines, and chloramphenicol, increasing their MICs (Figs. 4B-C). In some cases, although not enough to change the MIC, PYO and TOX had drug-antagonistic effects that could be measured by an increase in optical density at the pre-MIC concentration (see antibiotics marked with “+” in Figs. 4B-D). Examples include exposure to PYO together with tetracycline or sulfamethoxazole/trimethoprim (Fig. 4D). Similar antagonistic effects were observed for cells exposed to TOX and levofloxacin (Fig. 4D). On the other hand, PYO and TOX acted synergistically with other drugs, potentiating their toxicity, resulting in lower MICs or lower optical densities at the pre-MIC concentrations when the metabolites were present. This was the case for tobramycin and meropenem in the presence of PYO (Figs. 4B, 4D), and for sulfamethoxazole/trimethoprim, tobramycin, meropenem, and ceftazidime in the presence of TOX (Figs. 4C-D). Cases where the synergistic effect was only visible at the pre-MIC concentrations are marked as “-” in Figs. 4B-D. We also tested the effect of PYO and TOX on colistin susceptibility, with both enhancing its toxicity. In the absence of these metabolites, *B. multivorans* 1 was completely resistant to colistin at all the concentrations tested (MIC > 4.096 mg/mL, Table S4), and therefore colistin is not included in Figs. 4B-C. However, addition of PYO caused a decrease in optical density (Fig. S11C), and TOX caused a dramatic drop in the MIC (Table S4), indicating that these metabolites can act synergistically with polymyxins (Meirelles et al., 2021; Schiessl et al., 2019; Zhu et al., 2019). Overall, these results show that the effects of redox-active secondary metabolites on antibiotic susceptibility can be dramatic and are distinct for different classes of drugs.

**Figure 4.**
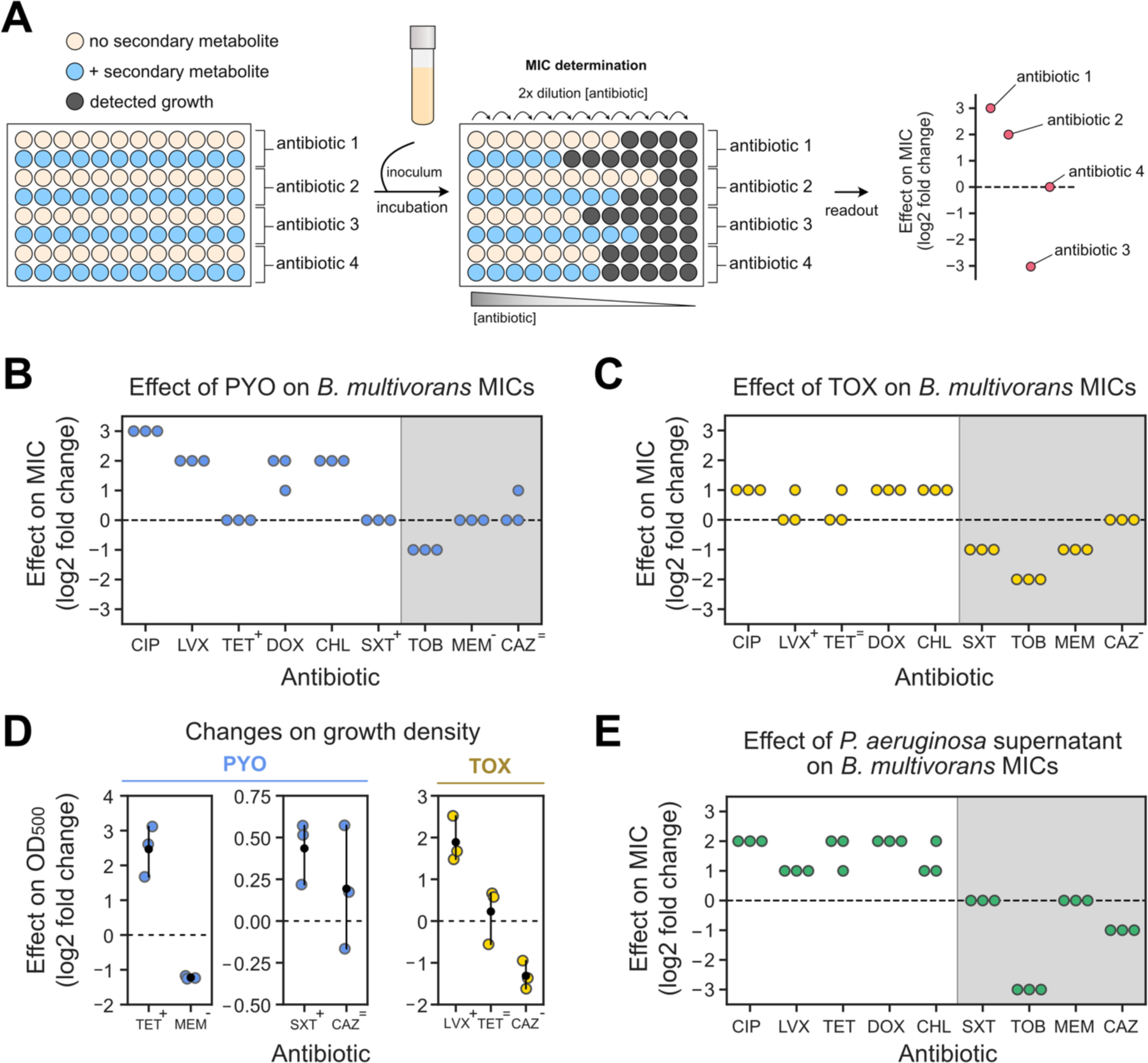
Assessing the effects of secondary metabolites on antimicrobial susceptibility testing. **A.** Experimental design used during MIC tests that account for the effect of secondary metabolites on resistance levels. **B.** Effects of PYO (100 µM) on *B. multivorans* 1 MICs (for each antibiotic, n = 3). **C.** Effects of TOX (50 µM) on *B. multivorans* 1 MICs (for each antibiotic, n = 3). In B and C, symbols above the antibiotic names represent effects of PYO and TOX on growth density displayed in D (“+” represents increase in density, “-” represents decrease in density, and “=” represents no consistent change in density). **D.** Effects of PYO (two plots on the left) and TOX (right) on the growth density at the pre-MIC antibiotic concentrations during MIC assays. Antibiotics shown are the ones previously highlighted by the symbols in panels B and C. Note that scales are different for each plot. For normalized absorbance values, see Fig. S11A-B. **E.** Effects of *P. aeruginosa* WT supernatant (i.e. PYO present) on *B. multivorans* 1 MICs (for each antibiotic, n = 3). For experimental design, see Fig. S10B. Grey shading in B, C and E represent antibiotics for which metabolite-mediated increase in resilience was not observed under the studied conditions. In panels D, the black dots mark the means and error bars represent 95% confidence intervals. CIP, ciprofloxacin; LVX, levofloxacin; TET, tetracycline, DOX, doxycycline, CHL, chloramphenicol, SXT, sulfamethoxazole/trimethoprim; TOB, tobramycin; MEM, meropenem; CAZ, ceftazidime.

Recognizing that these effects might be generalizable for a wide range of unknown molecules for which purified compounds are unavailable, we sought to determine whether modified MIC protocols could be agnostic to any specific metabolite secreted by a pathogen. Accordingly, we used spent media from grown cultures (i.e., containing secondary metabolites) mixed with fresh media (Fig. S10B and Materials and Methods for experimental design) to modify the traditional MIC assay. As proof of principle, we tested the susceptibility of our model organism, *B. multivorans* 1, to all the drugs mentioned above, using spent medium from *P. aeruginosa* WT (Fig. 4E). We predicted that the phenazine-producing *P. aeruginosa* cultures would change the MIC like exogenously added PYO did in the previous experiments. As expected, *P. aeruginosa* spent medium increased the MIC of *B. multivorans* against fluoroquinolones, tetracyclines, and chloramphenicol (Fig. 4E). We did not see changes in MIC against sulfamethoxazole/trimethoprim (Fig. 4E), likely because our experimental design led to a four-fold dilution of the active metabolite(s) from its initial concentration in the spent medium. For instance, the PYO concentration was ∼80-100 µM in the original culture from which the spent medium was taken, but only ∼20-25 µM in the final assay (see Materials and Methods). Because the impact of exogenously adding 100 µM PYO was already small (Fig. 4D), it is not surprising that we observed no effect using the diluted spent media. Still, in agreement with our experiments using purified PYO, *P. aeruginosa* spent medium potentiated the efficacy of tobramycin, reducing the MIC (Fig. 4E). On the other hand, unlike our findings with purified PYO, *P. aeruginosa* spent medium decreased the MIC against ceftazidime (Fig. 4E). To evaluate if the results observed for *P. aeruginosa* spent medium were caused by phenazines, we also tested spent medium from the *P. aeruginosa* Δ*phz* mutant (Fig. S11D). We saw no significant increase in the *B. multivorans* MICs when Δ*phz* spent medium was used (Fig. S11D), suggesting that the antagonistic effects against these drugs were due to phenazines. Finally, we also tested the impact of spent medium from *B. multivorans* on its own MICs (Fig. S11E). We saw no effect on MICs for fluoroquinolones or sulfamethoxazole/trimethoprim, but we did see changes for the other drugs: *B. multivorans* spent medium increased MICs against tetracyclines, chloramphenicol, and meropenem but decreased MICs against tobramycin and ceftazidime (Fig. S11E). Altogether, our results suggest that metabolites secreted by opportunistic pathogens can substantially alter MIC levels in ways that are overlooked by current protocols.

## DISCUSSION

Redox-active secondary metabolites, produced by organisms throughout the tree of life (Jacob et al., 2011), are of particular interest due to their multifaceted and nuanced effects, which depend on the environmental and physiological conditions experienced by the cells when exposed to them (Glasser et al., 2017; Meirelles and Newman, 2018). Within the clinical context, their dynamic functions range from the support of biofilm development to serving as virulence factors that are toxic to host cells (Dietrich et al., 2013; Lau et al., 2004; Liu and Nizet, 2009; Ramos et al., 2010; Saunders et al., 2020). In this study, we found that these molecules can also modulate antibiotic resilience, altering susceptibility levels to drugs commonly used to treat infections. Together with recent evidence demonstrating that these molecules can increase mutation rates to antibiotic resistance against certain drugs (Meirelles et al., 2021), our results point towards the production of redox-active secondary metabolites by opportunistic pathogens as an underappreciated route for the evolution of antimicrobial resistance.

To better predict contexts where redox-active secondary metabolites might affect antimicrobial susceptibility, it is necessary to understand the molecular mechanisms involved in the process. Our results support the role of SoxR as a broad redox sensor in nature (Dietrich et al., 2008). This transcription factor has been primarily studied in enteric bacteria (*E. coli* and *Salmonella* sp.) and in *P. aeruginosa*, where it senses redox-active molecules through oxidation of its Fe-S cluster (Dietrich et al., 2006; Gu and Imlay, 2011; Lee et al., 2015; Sheplock et al., 2013; Singh et al., 2013). However, SoxR is widely distributed throughout the bacterial domain, with homologs enriched in the *Actinobacteria* and *Proteobacteria* (Glasser et al., 2017). Many of these organisms include well-established or emerging opportunistic pathogens, such as species within the genus *Mycobacterium*, *Nocardia*, *Burkholderia* (including the Bcc species studied here), *Ralstonia*, *Acinetobacter*, and *Stenotrophomonas*, among others (Dietrich et al., 2008; Glasser et al., 2017). However, we know little about the redox-active molecules SoxR might sense and the responses it might control in these organisms. Our work demonstrates that, if focused on antibiotic susceptibility, special attention should be given to SoxR-regulated efflux systems and to the drugs these systems might be able to transport. SoxR homologs in other pathogens might control the transport of yet-to-be-discovered redox-active secondary metabolites that affect antibiotic susceptibility, whether the metabolite is produced endogenously or by a neighbor species.

While we have shown that certain secondary metabolites can dramatically modulate antibiotic susceptibility *in vitro*, it is important to recognize that these effects have yet to be confirmed *in vivo*. Testing the relevance of this phenomenon in the host context is an essential next step, and we hope our study will stimulate future work on this topic. Moreover, there is still a long path ahead when considering interspecies interactions during infections, mostly due to our still limited understanding of the colonization dynamics in polymicrobial infections. This is true even for CF, a well-studied infection context. Although we know certain species can be found infecting patients at the same time, our understanding of the frequency of these interactions is much less clear. For example, *P. aeruginosa* can be found with *Staphylococcus* or Bcc (Chmiel et al., 2014; Folescu et al., 2015; Lipuma, 2010; Schwab et al., 2014), but the prevalence of such co-infections among patients is not well documented. Similarly, *B. gladioli* is among the most common *Burkholderia* species isolated from CF sputum (Lipuma, 2010), but we do not know which other microbes typically co-reside with this species. Moreover, how these organisms co-exist within infections is poorly constrained due to very limited data on their spatial distribution within patients, though methods for accessing this are improving (DePas et al., 2016; Earle et al., 2015; Shi et al., 2020; Wilbert et al., 2020). While our study suggests that redox-active metabolites such as PYO and TOX may be broad modulators of antibiotic susceptibility, to accurately predict how/which neighboring species might be affected by such molecules, future work must characterize community dynamics and spatial proximity between co-infecting organisms within individual patients.

Our results also highlight the need for redesign of antimicrobial susceptibility tests to make them more accurately mimic the context of infections. While there is room for improvement across a range of parameters, inclusion of secondary metabolites from spent supernatants of infecting strains in AST protocols could be a simple and constructive first step, even without knowledge of the identity of the metabolite(s) that impact the results. Because the vast majority of secondary metabolites are still uncharacterized (Skinnider et al., 2020), it is important to use phenotypic screens to identify the existence of an unknown whose identity can later be determined through follow-up research. Our case study using PYO and TOX as models for evaluating the impact of redox-active secondary metabolites on antibiotic resilience serves as a proof of principle that awareness and understanding of these interactions has the potential to inform drug selection for more effective treatment. For example, our results suggest that when in the presence of *P. aeruginosa*, treating Bcc with fluoroquinolones, tetracyclines, chloramphenicol, or sulfamethoxazole/trimethoprim would likely be less effective, increasing the chances of the evolution of resistance (Meirelles et al., 2021). For these situations, drugs like meropenem or ceftazidime would possibly be more appropriate. TOX-producing *B. gladioli* might also significantly impact Bcc susceptibility to antibiotics, mainly decreasing the efficacy of fluoroquinolones. Finally, *B. multivorans* appears to produce unknown metabolites that reduce its susceptibility to certain drugs, such as tetracyclines, chloramphenicol, and meropenem, which could be investigated in more detail in the future.

More generally, we end by observing that approaches that have been employed to identify polymicrobial and/or microbe-host interactions mediated by secondary metabolites in the context of the gut microbiome (Agus et al., 2021; Vernocchi et al., 2016) can and should be leveraged more strongly in the context of infectious disease. While some studies have begun to light the way (Bauermeister et al., 2021; Garg et al., 2017), there remains tremendous potential for discovery of secondary metabolite-mediated effects that shape clinical outcomes. To identify endogenously produced molecules that impact antimicrobial susceptibility, we must ask: Who is there? What are they making? What are the responses induced in the community by the presence of specific molecules? And, finally, which drugs are affected by these responses? High-throughput methods combined with various “systems approaches” used for studying how drugs interact with the gut microbiome (Brochado et al., 2018; Maier et al., 2021, 2018; Zimmermann et al., 2021) could be adapted to study how metabolites produced by pathogens in different infection contexts might affect their interactions with each other and their susceptibility to antibiotics. We anticipate that our findings with PYO and TOX are likely just the tip of the iceberg, and we hope growing awareness of the potential ubiquity of these type of interactions will enhance our understanding of and ability to control polymicrobial infections.

## MATERIALS AND METHODS

### Media and incubation conditions

Most of the media and conditions used followed previous descriptions (Meirelles et al., 2021). The defined medium mostly used in this work was the glucose minimal medium (GMM), comprising 20 mM glucose (or 10 mM, if specified), 50 mM KH_2_PO_4_/K_2_HPO_4_ (pH 7.2), 42.8 mM NaCl, 9.35 mM NH_4_Cl, 1 mM MgSO_4_, 1x MEM Amino Acids (AA) solution (MilliporeSigma, Cat. No. M5550), and a trace elements solution (Widdel and Pfennig, 1981). As described previously (Meirelles et al., 2021), the medium was prepared by autoclaving all the components together, except for the glucose and the 1,000x trace elements stock solution, which were sterilized through filtration and added after the autoclave step. Autoclave step proceeded for 20 minutes at 121°C. As previously noted (Meirelles et al., 2021), autoclaving MgSO_4_ together with other media components is essential for consistent production of PYO by our WT *P. aeruginosa* UCBPP-PA14 strain in this medium. Two other media were used: Luria–Bertani (LB) Miller broth (BD Biosciences) and BBL cation-adjusted Mueller–Hinton II (MH) broth (BD Biosciences). Their preparation followed the manufacturer’s instructions. Importantly, for the MH broth medium, the autoclave step was 10 min. 1.5% Bacto agar (BD Biosciences) was used for solid media unless mentioned otherwise (see strains construction section below). Pyocyanin (PYO) was synthesized and purified following published protocols (Cheluvappa, 2014; Costa et al., 2017), and 10 mM stock solutions were prepared using 20 mM HCl and stored at −20°C. Toxoflavin (TOX) (MedChemExpress) was dissolved in dimethyl sulfoxide (DMSO) to make 10 mM stock solutions that were prepared on the same day of the experiment. Phenazine-1-carboxylic acid (PCA) (Princeton Bio) was dissolved in 20 mM NaOH and stored at −20°C. Finally, experiments using PYO, TOX or PCA always had negative controls with equivalent volumes of the solvents used (20 mM HCl, DMSO, or 20 mM NaOH, respectively). Unless otherwise specified, incubations were performed at 37°C, with shaking for liquid cultures at 250 rpm.

### Strain construction

We performed genetics in the *B. multivorans* AU42096 strain, our model Bcc organism. We refer to this strain as *B. multivorans* 1 (Meirelles et al., 2021) (Table S1). An unmarked deletion of the Bmul_3929 gene (*soxR* homolog) was made through homologous recombination using the pEX18Tc plasmid (Hoang et al., 1998; Huang and Wilks, 2017). Briefly, ∼800 bp fragments upstream and downstream of the gene were amplified and cloned into the pEX18Tc suicide vector using Gibson assembly (Gibson et al., 2009). Amplifications were done using *B. multivorans* 1 genomic DNA (gDNA, extracted with DNeasy Blood & Tissue Kit, Qiagen), cleaned up using the Monarch PCR Purification kit (New England Biolabs), and used in a Gibson assembly reaction (New England Biolabs) together with a PCR-amplified fragment containing the entire pEX18Tc sequence. The assembled construct was transformed into *E. coli* S17 (Simon et al., 1983), with selection in LB with 10 µg/mL tetracycline and incubation at 30°C. The assembled plasmid was identified by colony PCA, verified by Sanger sequencing (Laragen), and inserted into *B. multivorans* 1 genome by biparental conjugation using a modified version of previously published protocols (Choi and Schweizer, 2006; Dubarry et al., 2010). Briefly, *E. coli* S17 (containing the pEX18Tc-based deletion plasmid) and the *B. multivorans* 1 WT strain were grown overnight in LB (with 10 µg/mL tetracycline for the *E. coli* strain) and then diluted in the same medium to OD_600_ of 0.05 and 0.025, respectively. Cultures were grown until OD_600_ of ∼0.5; if one of the cultures reached the final OD earlier, this culture was moved to a non-shaking incubator (also 37°C) until the other was ready for the next steps. When ready, 400 µL of each culture were mixed into a Falcon tube and left stand (non-shaking) for 1 hour. Next, 100-200 µL of the mixed cultures were plated on top of polycarbonate membranes (MilliporeSigma, Cat. No. WHA10417006) on LB plates, followed by an incubation step of ∼15 hours at 37°C. Next, cells were scrapped from the filters, resuspended in 0.9% NaCl, plated on to VBMM agar plates (3 g/L trisodium citrate, 2 g/L citric acid, 10 g/L K_2_HPO_4_, 3.5 g/L NaNH_4_PO_4_·4H_2_O, 1 mM MgSO_4_, 100 μM CaCl_2_, pH 7) containing 100 µg/mL of tetracycline (Choi and Schweizer, 2006), and incubated for 24-48 hours at 30°C. This resulted in colonies of *B. multivorans* 1 that were merodiploids containing the construct integrated in their genomes, which were then plated on to LB containing 10% sucrose. Importantly, LB plates lacked NaCl and contained only 1.05% agar. This was relevant because we noticed that even though pEX18Tc contains a *sacB* cassette, the *sacB* counter selection was not effective in our *B. multivorans* 1 strain, as has been observed for other *Burkholderia* strains (Barrett et al., 2008). However, these conditions still allowed the screening of colonies that lost the integrated plasmid. We noticed that such colonies presented a spreading “flat” morphology under these conditions, while the merodiploids were round “thick” colonies (Fig. S12). The flat colonies were rare (roughly 1 in ∼200 colonies) and, as expected, ∼50% of them were WT genotype and ∼50% were unmarked deletions. These were screened by PCR and verified by both Sanger sequencing and whole-genome sequencing.

Complementation of the Δ*soxR* mutant was also done through homologous recombination with the re-insertion of the gene in its native site following the same protocols described above. The only difference was that, in this case, a unique fragment from ∼800 bp fragments upstream to ∼800 downstream region of *soxR* (i.e., containing the *soxR* gene) was amplified from *B. multivorans* 1 WT gDNA and cloned into the pEX18Tc. This was then inserted in the Δ*soxR* mutant through homologous recombination was described above, with PCR and whole-genome sequencing verifications. Information on all primers and strains used is available in Table S1.

### Whole-genome sequencing and draft genome assembly and annotation

Genomic DNA (gDNA) was extracted from the following strains using the DNeasy Blood & Tissue kit (Qiagen): *B. multivorans* 1 WT; *B. multivorans* 1 Δ*soxR*; *B. multivorans* 1 Δ*soxR* comp (complementation of *soxR* in the Δ*soxR* background). Library preparation was performed by the Microbial Genome Sequencing Center (MiGS) (Pittsburgh, Pennsylvania, USA) and included (i) 2 x 150 bp paired-end Illumina sequencing for all these strains (minimum of 400-Mb sequencing output per sample, with approximately 50-60x coverage), and (ii) an additional Nanopore sequencing for the *B. multivorans* 1 WT strain (minimum of 300 Mb Long Reads) for draft genome assembly and annotation of this strain.

Draft genome assembly and annotation of the *B. multivorans* 1 WT strain was performed by the MiGS analysis pipeline, and included: (i) quality control and adapter trimming performed using bcl2fastq (for Illumina reads) and porechop (for Nanopore reads, available at: https://github.com/rrwick/Porechop); (ii) assembly (using both Illumina and Nanopore reads) performed using Unicycler (Wick et al., 2017); (iii) assembly statistics assessed using QUAST (Gurevich et al., 2013), which can be found in the “BM1_assembly_metrics.tsv” file in the supplementary material; and (iv) draft assembly annotation performed using Prokka (Seemann, 2014). The assembly and annotation files can also be found in the supplementary material as “BM1_assembly.fasta” and “BM1_annotation.gff”, together with a summary of the assembled contigs “BM1_contigs_summary.tsv”.

To check for the Δ*soxR* and Δ*soxR* comp strains, quality control was performed using Trimmomatic (version 0.39) (Bolger et al., 2014) with the following settings: LEADING:27 TRAILING:27 SLIDINGWINDOW:4:20 MINLEN:35. Mutations were then identified using breseq (version 0.35.7) (Deatherage and Barrick, 2014) using the draft annotation we described above. This was important because we observed dozens of potential non-related mutations that appeared in the genome during the process of making these strains. This was not surprising since *B. multivorans* 1 is a clinical strain that has not been extensively used in the laboratory, with the exception of our previous work that did not include genetic manipulation (Meirelles et al., 2021). Despite these additional potential mutations, we were able to confirm the correct *soxR* deletion and complementation, which resulted in the phenotypes described in our results. Whole-genome sequence data for the strains studied was submitted to the NCBI Sequence Read Archive under the accession number TBD.

### Assessment of the presence of SoxR/RND-9 locus in the studied strains

The SoxR/RND-9 locus presence was determined by three complementary methods. First, we directly inspected the genomes of the *Burkholderia* species studied that are available at the *Burkholderia* Genome Database (BGD) (Winsor et al., 2008) and Integrated Microbial Genomes and Microbiomes IMG/M (Chen et al., 2021; Mukherjee et al., 2021). These included *B. cenocepacia* J2315, *B. cenocepacia* K56-2, *B. cepacia* ATCC 25416, and *B. glumae* 336gr-1. We also inspected the draft genomes for the strains we performed whole-genome sequencing, which included all the genotypes of our *B. multivorans* 1 strain. Second, we performed PCR amplification attempts for a fragment containing part of the SoxR/RND-9 genome locus in all strains. Primers were designed using consensus sequences of the region known from the available genomes (see primers in Table S1). As positive controls for this screen, we used primers for 16S rRNA amplification. For all PCRs, overnight cultures for all strains were grown in LB and 1 µL of the cultures were used during amplification attempts. Finally, we also checked annotated genome of closely-related strains available at the BGD or IMG/M. We could not find strains of *B. glumae* or *B. gladioli* that had the SoxR/RND-9 genome locus.

### Phylogenetically analyses

Phylogenetic analysis included 15 *Burkholderia* strains for which 16S sequences were either generated by Sanger sequencing (Laragen; primers available in Table S1) or retrieved from the BGD database or GenBank. For generated sequences, contigs were prepared using MacVector (version 18.2.0). Sequences were aligned with MAFFT (version 7.490, alignment available in the supplementary material) (Katoh et al., 2002; Katoh and Standley, 2013), and phylogeny was reconstructed through Bayesian inference using MrBayes (version 3.2.7) (Ronquist et al., 2012). Briefly, GTR was used as initial model for two independent analyses, each consisting of three heated chains and one cold chain. Markov Chain Monte Carlo sampling ran for one million generations (additional parameters: diagnfreq=1000; samplefreq=100; relburnin=yes; and burninfrac=0.25), which was enough to reach convergency (i.e. average standard deviation of split frequencies < 0.01). The final tree was edited in FigTree (version 1.4.4, available at: https://github.com/rambaut/figtree/releases) and polished in Affinity Designer (Serif, version 1.10.4).

### RNA-seq experiment and data analysis

Four different conditions were prepared for the RNA-seq experiment: (i) *B. multivorans* (no PYO added); (ii) *B. multivorans* + 100 µM PYO; (iii) *B. multivorans* + *P. aeruginosa* WT (co-culture, PYO and other phenazines produced); and (iv) *B. multivorans* + *P. aeruginosa* Δ*phz* (co-culture, no PYO or phenazines produced). For these experiments, we used our *B. multivorans* 1 WT strain. From freshly streaked LB plates (< 2 days old), overnight cultures of *B. multivorans*, *P. aeruginosa* WT and *P. aeruginosa* Δ*phz* were grown in GMM (20 mM glucose) + AA. Cells were washed (12500 rpm for 2 min) and resuspended in the same medium. OD_500_ values were measured and adjusted for the start of the experiment. Due to growth differences, the conditions involving co-cultures started with four times more *Burkholderia* cells than *Pseudomonas* cell. ODs used were as follows: conditions (i) and (ii) had initial OD_500_ = 0.04 (only *Burkholderia* was present); conditions (iii) and (iv) had final OD_500_ = 0.05, composed from 0.04 of *Burkholderia* cells and 0.01 of *Pseudomonas* cells. Cultures were prepared in 7 mL media using GMM (20 mM glucose) + AA, with either 100 µM PYO (used in condition 2) or the respective amount of HCl added to the cultures. The four tubes were incubated for ∼9 hours, and then collected for RNA extraction. For this, 0.7 mL of cultures were spun down (14000 rpm for 2 min), supernatants were removed, and the pellets were immediately frozen using liquid nitrogen and stored at -80°C. At this same time, PYO concentration produced by the co-culture containing the WT *P. aeruginosa* was measured from the supernatant using absorbance at OD691 (Reszka et al., 2004), and ∼ 60 µM was detected at the time of sampling.

Next, RNA extraction was performed using the RNeasy kit (Qiagen) following the manufacturer’s instructions. Samples were thawed on ice and resuspended in TE buffer (30 mM Tris.Cl, 1 mM EDTA, pH 8.0) containing 15 mg/mL of lysozyme and proteinase K solution (20 mg/mL, Qiagen), followed by an incubation with vortex and lysis steps described in the kit. Purified RNA was then sent for sequencing at the MiGS Center as done for whole-genome sequencing. The facility pipeline included: (i) DNase treatment (RNAse free) (Invitrogen); (ii) library preparation using Qiagen FastSelect and Library Prep kits (Qiagen) and Ribo-Zero Plus kit (Illumina); (iii) sequencing using a NextSeq500, with 1 x 75 bp reads for the four conditions, with a minimum of 16M reads for conditions 1 and 2 (single *Burkholderia* cultures), or 24M reads for conditions 3 and 4 (co-cultures); (iv) demultiplexing and adaptors trimming using bcl2fastq (version 2.20.0.422). Next, low-quality bases were removed using Trimmomatic (version 0.39) with the following settings: LEADING: 27 TRAILING: 27 SLIDINGWINDOW: 4:20 MINLEN: 35 (Bolger et al., 2014). Genome mapping and calculation of number of reads per gene was performed through Rockhopper (version 2.03) (Tjaden, 2015) using the *Burkholderia multivorans* ATCC 17616 genome available in the software as reference. Mapping was done against all three chromosomes (NC_010084; NC_010086; NC_010087), as well as against the pBMUL01 and pTGL1 plasmids (NC_010070 and NC_010802, respectively). Settings within the software were: 0.15 for allowed mismatches; 0.33 for minimum seed length; 500 for max bases between paired reads; 0.5 for minimum expression of UTRs and ncRNAs; and the “strand specific” option was unmarked. Even though the library preparation pipeline included an rRNA depletion step, we retrieved a large amount of rRNA sequences, which were manually deleted from the read-count table exported by Rockhopper. Data exploration and analysis were performed using the online Degust tool (Powell, 2015). With Degust, counts per million (CPM) for each gene were used for sequencing depth normalization and log2 fold changes are presented for two different comparison. In comparison 1, condition 1 (*B. multivorans* - no PYO added) was used as a baseline control during fold-changes calculation detected for condition 2 (*B. multivorans* + 100 µM PYO). In comparison 2, condition 4 (*B. multivorans* + *P. aeruginosa* Δ*phz*) was used as baseline control during fold-changes calculation detected for condition 3 (*B. multivorans* + *P. aeruginosa* WT). See Table S2 for the full results. Because our samples consisted of one replicate, no statistical tests were performed. Instead, we used this RNA-seq data as a screen where specific genes of interest (particularly related to efflux) were next explored by qRT-PCR experiments. RNA-seq sequence data was submitted to the NCBI Sequence Read Archive under the accession number TBD.

### Quantitative reverse transcription PCR (qRT-PCR)

#### Experiment 1: validation of RNA-seq

The set up for this experiment was exactly the same as the RNA-seq described above, with the exception that three independent replicates were prepared. For each strain used in this experiment (*B. multivorans* 1, *P. aeruginosa* WT, *P. aeruginosa* Δ*phz*), three independent overnight cultures were grown in GMM (20 mM glucose) + AA. Each of overnights were inoculated from three different spots from the freshly streaked LB plates of each strain. Next, each one of these overnight cultures was used in the inoculum preparation of the four different conditions, as described in the RNA-seq section. Again, the conditions were: (i) *B. multivorans* (no PYO); (ii) *B. multivorans* + 100 µM PYO; (iii) *B. multivorans* + *P. aeruginosa* WT (co-culture, PYO and other phenazines produced); and (iv) *B. multivorans* + *P. aeruginosa* Δ*phz* (co-culture, no PYO or phenazines produced). A total of 12 tubes were prepared (three for each conditions). These were grown for ∼9 hours, after which the pellets were collected (from 0.7 mL of culture), immediately frozen using liquid nitrogen, and stored at -80°C for later RNA extraction.

### Experiment 2: measuring the SoxR effect on PYO-mediated induction of RND-9

Four different conditions were prepared for this qRT-PCR experiment: (i) *B. multivorans* Δ*soxR*; (ii) *B. multivorans* Δ*soxR* + 100 µM PYO; (iii) *B. multivorans* Δ*soxR* comp; (iii) *B. multivorans* Δ*soxR* comp + 100 µM PYO. Three independent overnight cultures of each strain were grown in GMM (20 mM glucose) + AA, cells were then washed and resuspended at an OD_500_ of 0.04 in fresh GMM (20 mM glucose, 7 mL culture). Three replicates were prepared for each condition, with a total of 12 samples. Depending on the condition, either 100 µM PYO or the respective amount of HCl was added to the cultures. Cultures were then incubated for ∼10 hours, after which the pellets were collected (from 0.7-1 mL of culture), immediately frozen using liquid nitrogen, and stored at -80°C for later RNA extraction.

### Experiment 3: measuring the SoxR effect on TOX-mediated induction of RND-9

Six different conditions were prepared for this qRT-PCR experiment. Each of the three tested strains (*B. multivorans* WT, Δ*soxR* and Δ*soxR* comp) were grown with and without 50 µM TOX. Three independent overnight cultures of each strain were grown in GMM (20 mM glucose) + AA, cells were then washed and resuspended at an OD_500_ of 0.05 in fresh GMM + AA (20 mM glucose, 5 mL culture). Again, three replicates were prepared for each condition, with a total of 18 samples. Either 50 µM TOX or the respective amount of DMSO was added to the cultures. These cultures were incubated for ∼20 hours, cells were pelleted (from 0.7 mL of culture), immediately frozen using liquid nitrogen, and stored at -80°C for later RNA extraction.

For the next steps, we followed previously published protocols (Babin et al., 2016; Meirelles et al., 2021; Meirelles and Newman, 2018). RNA extraction was performed as described in these studies and in the RNA-seq experiment using the RNeasy kit (Qiagen). Contaminant gDNA was removed using TURBO DNA-free kit (Invitrogen, Waltham, Massachusetts, USA), and cDNA was synthesized using the iScript cDNA Synthesis kit (Bio-Rad, Hercules, California, USA) (a total of 0.8 μg of total RNA was used), following the manufacturer’s instructions. Then, qRT-PCR reactions were performed using iTaq Universal SYBR Green Supermix (Bio-Rad) (total of 20 μL per reaction) using a 7500 Fast Real-Time PCR System machine (Applied Biosystems, Waltham, Massachusetts, USA). For additional details on the protocol, see Meirelles and Newman (2018). Finally, within each run, standard curves for each primer pair were prepared using known concentration of *B. multivorans* WT gDNA to calculate amounts of cDNA for each of the target genes. The gene Bmul_2161 (annotated as *uvrC*, coding for excinuclease ABC subunit C) was used as housekeeping gene during normalizations. As an additional control, we also ran reactions with the housekeeping gene Bmul_1456 (annotated as *rumA* gene, coding for the 23S rRNA 5-methyluridine methyltransferase) (Schnetterle et al., 2021). Primer pairs sequences are available in Table S1.

Data showing total *uvrC*-normalized cDNA levels and/or the log2 fold change in expression are shown in Fig. 1C-D, Fig. 2B, Fig. 3F, Fig. S1, Fig S2, and Fig. S9. Normalizations for cDNA measurement followed what we have previously described for *oprI*-normalized cDNA in *P. aeruginosa* (Meirelles et al., 2021) but using the *uvrC* gene instead. In brief, the cDNA estimated for a certain gene in a certain sample was divided by the respective cDNA estimated for *uvrC* in the same sample (Meirelles et al., 2021). In some occasions, we also present the data as fold changes. These were calculated for each replicate relative to the mean cDNA value of the replicates within the negative control. The “no treatment” samples (i.e. “no PYO” or “no TOX”) were used as negative controls for conditions where *B. multivorans* was in single-species cultures (in experiments 1, 2 and 3); meanwhile, the “*B. multivorans* + *P. aeruginosa* Δ*phz*” samples was used as negative control for the “*B. multivorans* + *P. aeruginosa* WT” condition (in experiment 1). Importantly, we noticed the one replicate for three following conditions had significantly lower amounts of cDNA by the end of our qRT-PCR protocol: “*B. multivorans* + *P. aeruginosa* WT” and “*B. multivorans* + *P. aeruginosa* Δ*phz*” within experiment 1, and “Δ*soxR* no TOX” in experiment 3. These were removed from the analysis as noted in the legends of figures where this data is displayed.

### Antibiotics used for tolerance and resistance assays

The following antibiotics were used in this study: ciprofloxacin (Fluka), levofloxacin (Sigma-Aldrich), tetracycline (tetracycline hydrochloride, Sigma-Aldrich), doxycycline (doxycycline hyclate, Sigma-Aldrich), tobramycin (TCI), colistin (colistin sulfate salt, Sigma-Aldrich), ceftazidime (TCI, containing ca. 10% Na2CO3), chloramphenicol (Sigma-Aldrich), meropenem (meropenem trihydrate, Sigma-Aldrich), trimethoprim (Sigma-Aldrich), and sulfamethoxazole (Sigma-Aldrich). Stock solutions of the antibiotics used were prepared in different solvents. Ciprofloxacin was dissolved in 20 mM HCl; levofloxacin, tetracycline, doxycycline, tobramycin, colistin and ceftazidime were dissolved in deionized water; chloramphenicol, meropenem and trimethoprim-sulfamethoxazole (mixed at 1:1 ratio) were dissolved in DMSO. Stock concentrations used were of 1 mg/mL for ciprofloxacin and levofloxacin, and 10 mg/mL for all other antibiotics.

### Antibiotic tolerance assays with single species

A growth curve assay was used to measure resistance levels to PYO, following a previously described protocol (Meirelles et al., 2021). Briefly, cells of the respective strain used were grown from a fresh plate into overnight cultures in GMM + AA with 20 mM glucose. Cells were then pelleted, washed, and resuspended in four independent cultures per treatment at an OD_500_ of 0.05 in GMM + AA, with 10 mM glucose. Treatments involved addition or not of the redox-active secondary metabolite (PYO, PCA or TOX, at the respective concentration displayed in the figures). These four independent replicates were prepared in 7 mL (for PYO and PCA) or 5 mL cultures (for TOX), always using 18 × 150 mm glass tubes, and incubated for around 20 hours. This was enough for cells to reach stationary phase. Next, each individual culture was then split into a “no antibiotic” negative control, or an “+ antibiotic” treatment as done before (Meirelles and Perry et al, 2021), using 2 mL of culture per treatment in plastic Falcon tubes (VWR, Cat. No. 352059). After antibiotic treatment was added, cultures were then incubated for four hours under shaking conditions. Finally, cultures were serially diluted using Minimum Phosphate Buffer (50 mM KH_2_PO_4_/K_2_HPO_4_, 42.8 mM NaCl, pH 7.2) and plated for CFUs using LB agar plates. These plates were then incubated at room temperature, with CFUs counted after 36-48 hours. To check for slow-growing/late-arising colonies, plates were also checked after 5-7 days. This protocol was used for all the strains mentioned in the figures containing antibiotic assays with single species cultures. These included: *B. multivorans* 1 (WT, Δ*soxR* or Δ*soxR* comp strains), and *B. glumae* (WT and Δ*toxA* strains). The antibiotic used was ciprofloxacin (at 10 µg/mL), as indicated in the figure legends.

### Growth curves for measuring resistance to PYO and data analysis

A growth curves assay was used to measure resistance levels to PYO, following conditions previously described (Meirelles et al., 2021). Briefly, the several *Burkholderia* strains listed in Figs 2E and S4 (also see Table S1 for additional details) were used in this experiment. Each strain was grown overnight from a fresh plate in GMM with 20 mM glucose + AA. Cells were then pelleted, washed, and resuspended at an OD_500_ = 0.05 using the same medium (this was the initial OD used in the experiment). The experiment was set up in 96-well plates, with different concentration of PYO being used (0, 10, 100, or 200 µM). Three wells were prepared for each treatment, and each well was considered an independent replicate. The total volume within each well contained 150 µL of culture and 70 µL of mineral on top (used for evaporation prevention). Incubation proceeded for 24 hours at 37°C under shaking conditions. A Spark 10M plate reader was used (Tecan), with absorbance readings at an OD_500_ every 15 minutes.

Data analysis was performed using a custom Python software containing tools for data processing/analysis/visualization designed for the datasets exported from our plate readers (available at: https://github.com/jciemniecki/dknlab_tools, version 1.1.0). Growth curves for all the strains and treatments can be seen in Fig. S5. Next, growth rates were determined using linear fits of the linear range of these growth curves (time range used: 3-8 hours for all strains except the slow growers *B. cenocepacia* J2315 and *B. gladioli* 3; for which a range of 3-15 hours was used). This was done using a linear regression function (also available at the same GitHub repository). Means of the growth rates were calculated for each strain/treatment based on the data from the three replicates. Finally, for each strain, these growth rate means obtained for treatments containing the different concentrations of PYO (10, 100 and 200 µM) were normalized (i.e. divided by) the growth rate means obtained for the “No PYO” control to calculate the “growth rate ratio” displayed in Fig. 2E.

### Antibiotic tolerance assays with co-cultures

Antibiotic tolerance assays using co-cultures were performed using a membrane-separated 12-well tissue plate cultures containing 0.1 μm pore PET membranes (VWR, Cat. No. 10769–226), following previously protocols (Meirelles et al., 2021). These were used (i) to study how *P. aeruginosa* can have affect antibiotic tolerance levels of different strains *B. multivorans* 1; (ii) to study how multiple TOX producers (*B. glumae* and *B. gladioli* strains) can affect antibiotic tolerance levels of several Bcc species. Overnight cultures were grown for each strain in GMM (20 mM glucose) + AA, cells were pelleted, washed, and resuspended in the same medium at different OD500: 0.05 for redox-active metabolites producers; and 0.025 for the Bcc species (in which antibiotic susceptibility was evaluated). Producers (*P. aeruginosa*, *B. glumae* and *B. gladioli* strains) were cultured in the bottom part of the well using a volume of 600 µL, while Bcc were cultured in the upper part of the well using a volume of 100 µL. The experimental design can be seen in Fig. 3G. When grown alone (i.e., no producer), the Bcc strain was added to both bottom and upper part of the well. Cells were grown under shaking conditions (175 rpm) for ∼20 hours at 37°C, with the plates placed within airtight plastic container. Several wet paper towels were used to maintain humidity. Next, ciprofloxacin was then added (10 µg/mL), and cultures were incubated for additional four hours. Within each plate, three replicates (wells) were used for “no antibiotic” (negative control) and three replicates for “+ ciprofloxacin” treatment. After incubation, the upper part of the co-cultures (containing the Bcc cells) was plated for CFUs on LB agar plates using serial dilution as described above.

In addition to the co-cultures grown on the membrane-separated 12-well tissue plate, we also performed a co-culture assay using 18 x 150 mm glass tubes (Fig. 1B, right). Here, *B. multivorans* 1 WT was grown together with *P. aeruginosa* WT or Δ*phz* in 7 mL cultures with a total starting OD_500_ of 0.05 (0.025 for each species). Four independent replicates were prepared, cultures were grown for 20 hours, treated with ciprofloxacin (10 µg/mL), and plated for CFUs as described before. Differently from the membrane-separated plates used before, the two species were mixed in this assay. However, *P. aeruginosa* and *B. multivorans* colonies display different morphologies and growth rates, allowing for visual distinction in the “no antibiotic” control treatment. Moreover, *P. aeruginosa* is orders of magnitude more susceptible to the drug than *B. multivorans* under the concentration used (10 µg/mL), allowing for easy counting of the *B. multivorans* colonies in the “+ ciprofloxacin” treatments (Fig. 1B).

### Determination of minimum inhibitory concentrations (MICs)

#### MIC assays using pure redox-active molecules

MICs were determined following the EUCAST standard clinical methods suggested for broth microdilution assays (EUCAST, 2003), with modifications to account for the effects of redox-active molecules (e.g., PYO or TOX). Briefly, *B. multivorans* 1 cultures was grown from a fresh LB plate into MH broth, and were then diluted 1:100 into three independent replicates (5 mL) and grown for 14-18 hours until stationary phase. These three independent replicates were individually resuspended and used as inocula in 2-fold dilution series assays using 96-well microtiter plates, the 10 antibiotics mentioned before, and the redox-active molecules. The final concentrations used for PYO and TOX were 100 µM and 50 µM, respectively; controls involved adding the equivalent amount of the solvents used to solubilize these molecules (20 mM HCl for PYO, or DMSO for TOX). One antibiotic was used per plate, and the plate design can be seen in Fig. S10A, with each well having a final volume of 100 μL. Appropriate “no antibiotic” and “no cells” controls were always prepared (Fig. S10A). After inoculation, the microtiter plates were sealed with a plastic film (to avoid evaporation), wrapped in aluminum foil, and incubated without shaking at 37°C for 18 hours. Microtiter plates were always incubated in a single layer. After incubation, the wells were assessed for growth (turbidity) using a BioTek Synergy 4 plate reader (BioTek) or a Spark 10M plate reader (Tecan). Growth assessment was also done by naked eye as suggested by the reading guide for broth microdilutions by EUCAST (version 3.0, January 1^st^, 2021), and used for the OD_500_ thresholding (see the data analysis section below).

### MIC assays using spent media from pathogens’ cultures

these assays followed the same design described above, with the exception that instead of using pure redox-active molecules, a filtered-sterilized spent media from stationary cultures was mixed to fresh media and used during 2-fold dilution series assays. In summary, cultures of the strains in which spent media were analyzed (*P. aeruginosa* WT, *P. aeruginosa* Δ*phz*, and *B. multivorans* WT) were grown in MH broth, diluted back 1:100 into 50 mL cultures (initial OD_500_ = ∼0.05), and grown for around 12 hours for metabolite production. Cells were then spun down (5000 rpm for 10 minutes, twice), and the supernatant containing the spent media was collected and filter-sterilized through a large bottle top filter (Millipore, Cat. No. SCGPS02RE). The filtered spent media samples were plated on LB agar for contamination verification. Then, spent media samples were mixed with fresh MH broth (1/4 of the final volume was spent media) during 2-fold dilution series assays using the antibiotics indicated in Fig. 4E and Fig. S11D-E. Importantly, spent medium from *P. aeruginosa* WT contained ∼80-100 µM PYO measured by absorbance at 691 nm, meaning that the final PYO concentration on microtiter plates were around 20-25 µM. Incubation and growth assessment protocol followed what was described above.

### Data analysis

after turbidity measurements by OD_500_ absorbance and inspection by naked eye, we adopted OD_500_ = 0.11 as a threshold value for the indication of growth in the microtiter plates. This means that this was the lowest absorbance where growth could be detected by naked eye; any reading value lower than this was determined “no growth”. This threshold was applied to the raw absorbance values, and no normalizations by the “no cells” controls wells were done at this stage. MIC values for each treatment tested are available in Table S4. Next, calculation of the molecules’ “Effect on MIC (log2 fold change)” in Figs. 4B-C were performed as follows: for each antibiotic, MIC values detected for treatments containing PYO or TOX were divided by the MIC detected in respective negative controls (where HCl or DMSO were added, respectively). The plate design was always the same, and individual ratios were calculated within each replicate (for each antibiotic, all replicates were prepared in parallel on the same day and plate – see Fig. S10A for design). These ratios were then log2-transformed and are shown in Figs. 4B-C. Finally, for MICs detected in experiments containing spent media (from the *P. aeruginosa* strains or from *B. multivorans*), the plates were prepared on different days from the original experiments with the pure molecules. However, because MIC values were consistent (i.e. HCl or DMSO did not seem to affect the MICs measured at the amounts used), the same negative controls used for ratios’ calculation in the experiments with pure molecules were also used for the ratios’ calculation in experiments with spent media. Specifically, the “No PYO” controls (i.e. where HCl was added) were used for this purpose. Again, these ratios were log2-transformed and are shown in Fig. 4E and Fig. S11D-E.

Finally, we realized that sometimes, even though the MIC detected itself did not change, treatments could alter the growth detected at the pre-MIC concentrations. For this reason, we have also included plots with the OD_500_ absorbance measurements and the log2 fold change for OD_500_ absorbances between different treatment. For this, the absorbances measurements were normalized by their respective “no cells” control wells within the same plate. Then, as done for the MICs values, the fold changes were determined by calculating the ratio for each treatment (PYO or TOX) by their respective negative control within each replicate (e.g. absorbance for condition “+PYO replicate 1 in TET” divided by “+HCl replicate 1 in TET”), and these values were log2 transformed.

This was only done for experiments using pure molecules, since these were performed in parallel (i.e. same day) with their respective solvent control. Both OD_500_ normalized absorbance measurements and their log2 fold change are shown (Fig. S11A-B and Fig. 4D, respectively).

### Measurements of PYO and TOX concentrations

PYO concentrations in cultures containing *P. aeruginosa* WT were estimated by absorbance of the culture supernatant at 691 nm (Reszka et al., 2004) using standard curves with the pure molecule. A similar approach was used for estimation of TOX concentrations within cultures containing *B. glumae* and *B. gladioli*. UV-Vis spectra (200-800 nm) were recorded using a spectrophotometer (Beckman Coulter DU 800) for solutions containing pure TOX (100, 50, 25 µM), and supernatants of overnight cultures (grown GMM, 20 mM glucose + AA) of the following strains: *B. glumae* WT, B*. gladioli* 1 and *B. gladioli* 2. Moreover, supernatants of *B. glumae* Δ*toxA* and *B. multivorans* 1 strains grown in the same medium were used as negative controls since these strains cannot make TOX. Concentrations estimations were done for each sample based on characteristic absorbance of the molecule at 393 nm (Chen et al., 2012; Kim et al., 2004), calibrated by the absorbance values obtained for the standards using the pure molecule. Because only oxidized TOX absorbs at 393 nm, after being taken out of the shaking incubator, cultures were immediately spun down for supernatant collection. Absorbance spectra within the 300-500 nm range for these samples are shown in Fig. S8.

### Data wrangling, analysis, and visualization

Data wrangling and analysis involved a combination of processing in (i) Python (version 3.8), using the Pandas (version 1.3.1) (McKinney, 2010; The pandas development team, 2020) and NumPy (version 1.20.3) (Harris et al., 2020) libraries; or in (ii) Microsoft Excel (version 16.39), as described throughout the Materials and Methods section. All the visualization presented in the manuscript was performed using Matplotlib (version 3.4.2) (Hunter, 2007) and Seaborn (version 0.11.1) (Waskom, 2021). 95% confidence intervals presented in the figures were estimated with Seaborn while plotting the respective data using 10,000 bootstraps. Plots legends and their display organization within each figure were adjusted using Affinity Designer (Serif, version 1.10.4). The same software was used for drawing all the illustrations shown in the manuscript.

## Supporting information

Additional supplementary data - 16S alignment, draft genome assembly and annotation

## ACKNOWLEDGEMENTS

We thank Newman lab members for feedback and advice throughout the development of this project. In particular, we thank Megan Bergkessel for assistance and discussions about the RNA-seq, John Ciemniecki for developing and sharing the “dknlab_tools” package used during plate reader data analysis, John LiPuma (University of Michigan, CFF *Burkholderia cepacia* Research Laboratory and Repository) for providing the clinical *Burkholderia* strains, and the Microbial Genome Sequencing Center (MiGS) at Pittsburgh for sequencing of the samples. Finally, we thank Jong Ham and Inderjit Barphagha (Louisiana State University) for providing the *B. glumae* 336gr-1 strains (WT and Δ*toxA*), and Joanna Goldberg (Emory University) for providing the pEX18Tc plasmid. This work was supported by grants to D.K.N from the NIH (1R01AI127850-01A1, 1R01HL152190-01), and the Doren Family Foundation.

## AUTHORS’ CONTRIBUTIONS

LAM: Conceptualization, Methodology, Validation, Formal analysis, Investigation, Data curation, Writing – original draft preparation, Writing – review & editing, Visualization, Project administration. DKN: Conceptualization, Resources, Writing – review & editing, Supervision; Project administration; Funding acquisition.

## REFERENCES

1. Agus, A., Clément, K., Sokol, H., 2021. Gut microbiota-derived metabolites as central regulators in metabolic disorders. Gut 70, 1174–1182. doi:10.1136/gutjnl-2020-323071

2. Babin, B.M., Bergkessel, M., Sweredoski, M.J., Moradian, A., Hess, S., Newman, D.K., Tirrell, D.A., 2016. SutA is a bacterial transcription factor expressed during slow growth in *Pseudomonas aeruginosa*. Proc. Natl. Acad. Sci. USA 113, E597–605. doi:10.1073/pnas.1514412113

3. Balaban, N.Q., Helaine, S., Lewis, K., Ackermann, M., Aldridge, B., Andersson, D.I., Brynildsen, M.P., Bumann, D., Camilli, A., Collins, J.J., Dehio, C., Fortune, S., Ghigo, J.-M., Hardt, W.-D., Harms, A., Heinemann, M., Hung, D.T., Jenal, U., Levin, B.R., Michiels, J., Storz, G., Tan, M.-W., Tenson, T., Van Melderen, L., Zinkernagel, A., 2019. Definitions and guidelines for research on antibiotic persistence. Nat. Rev. Microbiol. 17, 441–448. doi:10.1038/s41579-019-0196-3

4. Barrett, A.R., Kang, Y., Inamasu, K.S., Son, M.S., Vukovich, J.M., Hoang, T.T., 2008. Genetic tools for allelic replacement in *Burkholderia* species. Appl. Environ. Microbiol. 74, 4498–4508. doi:10.1128/AEM.00531-08

5. Bartell, J.A., Blazier, A.S., Yen, P., Thøgersen, J.C., Jelsbak, L., Goldberg, J.B., Papin, J.A., 2017. Reconstruction of the metabolic network of *Pseudomonas aeruginosa* to interrogate virulence factor synthesis. Nat. Commun. 8, 14631. doi:10.1038/ncomms14631

6. Bauermeister, A., Mannochio-Russo, H., Costa-Lotufo, L.V., Jarmusch, A.K., Dorrestein, P.C., 2021. Mass spectrometry-based metabolomics in microbiome investigations. Nat. Rev. Microbiol. doi:10.1038/s41579-021-00621-9

7. Blair, J.M.A., Webber, M.A., Baylay, A.J., Ogbolu, D.O., Piddock, L.J.V., 2015. Molecular mechanisms of antibiotic resistance. Nat. Rev. Microbiol. 13, 42–51. doi:10.1038/nrmicro3380

8. Bolger, A.M., Lohse, M., Usadel, B., 2014. Trimmomatic: a flexible trimmer for Illumina sequence data. Bioinformatics 30, 2114–2120. doi:10.1093/bioinformatics/btu170

9. Brauner, A., Fridman, O., Gefen, O., Balaban, N.Q., 2016. Distinguishing between resistance, tolerance and persistence to antibiotic treatment. Nat. Rev. Microbiol. 14, 320–330. doi:10.1038/nrmicro.2016.34

10. Brochado, A.R., Telzerow, A., Bobonis, J., Banzhaf, M., Mateus, A., Selkrig, J., Huth, E., Bassler, S., Zamarreño Beas, J., Zietek, M., Ng, N., Foerster, S., Ezraty, B., Py, B., Barras, F., Savitski, M.M., Bork, P., Göttig, S., Typas, A., 2018. Species-specific activity of antibacterial drug combinations. Nature 559, 259–263. doi:10.1038/s41586-018-0278-9

11. Cheluvappa, R., 2014. Standardized chemical synthesis of *Pseudomonas aeruginosa* pyocyanin. MethodsX 1, 67–73. doi:10.1016/j.mex.2014.07.001

12. Chen, I.-M.A., Chu, K., Palaniappan, K., Ratner, A., Huang, J., Huntemann, M., Hajek, P., Ritter, S., Varghese, N., Seshadri, R., Roux, S., Woyke, T., Eloe-Fadrosh, E.A., Ivanova, N.N., Kyrpides, N.C., 2021. The IMG/M data management and analysis system v.6.0: new tools and advanced capabilities. Nucleic Acids Res. 49, D751–D763. doi:10.1093/nar/gkaa939

13. Chen, R., Barphagha, I.K., Karki, H.S., Ham, J.H., 2012. Dissection of quorum-sensing genes in *Burkholderia glumae* reveals non-canonical regulation and the new regulatory gene *tofM* for toxoflavin production. PLoS One 7, e52150. doi:10.1371/journal.pone.0052150

14. Chmiel, J.F., Aksamit, T.R., Chotirmall, S.H., Dasenbrook, E.C., Elborn, J.S., LiPuma, J.J., Ranganathan, S.C., Waters, V.J., Ratjen, F.A., 2014. Antibiotic management of lung infections in cystic fibrosis. I. The microbiome, methicillin-resistant *Staphylococcus aureus*, gram-negative bacteria, and multiple infections. Annals of the American Thoracic Society 11, 1120–1129. doi:10.1513/AnnalsATS.201402-050AS

15. Choi, K.-H., Schweizer, H.P., 2006. mini-Tn7 insertion in bacteria with single *att*Tn7 sites: example *Pseudomonas aeruginosa*. Nat. Protoc. 1, 153–161. doi:10.1038/nprot.2006.24

16. Costa, K.C., Glasser, N.R., Conway, S.J., Newman, D.K., 2017. Pyocyanin degradation by a tautomerizing demethylase inhibits *Pseudomonas aeruginosa* biofilms. Science 355, 170–173. doi:10.1126/science.aag3180

17. Coutinho, C.P., Dos Santos, S.C., Madeira, A., Mira, N.P., Moreira, A.S., Sá-Correia, I., 2011. Long-term colonization of the cystic fibrosis lung by *Burkholderia cepacia* complex bacteria: epidemiology, clonal variation, and genome-wide expression alterations. Front. Cell Infect. Microbiol. 1, 12. doi:10.3389/fcimb.2011.00012

18. Cruickshank, C.N., Lowbury, E.J., 1953. The effect of pyocyanin on human skin cells and leucocytes. Br J Exp Pathol 34, 583–587.

19. Davies, J., 2013. Specialized microbial metabolites: functions and origins. J Antibiot 66, 361– 364. doi:10.1038/ja.2013.61

20. Davies, J., Davies, D., 2010. Origins and evolution of antibiotic resistance. Microbiol. Mol. Biol. Rev. 74, 417–433. doi:10.1128/MMBR.00016-10

21. Deatherage, D.E., Barrick, J.E., 2014. Identification of mutations in laboratory-evolved microbes from next-generation sequencing data using breseq. Methods Mol. Biol. 1151, 165–188. doi:10.1007/978-1-4939-0554-6_12

22. DePas, W.H., Starwalt-Lee, R., Van Sambeek, L., Ravindra Kumar, S., Gradinaru, V., Newman, D.K., 2016. Exposing the three-dimensional biogeography and metabolic states of pathogens in cystic fibrosis sputum via hydrogel embedding, clearing, and rRNA labeling. MBio 7, e00796–16. doi:10.1128/mBio.00796-16

23. Depoorter, E., Bull, M.J., Peeters, C., Coenye, T., Vandamme, P., Mahenthiralingam, E., 2016. *Burkholderia*: an update on taxonomy and biotechnological potential as antibiotic producers. Appl. Microbiol. Biotechnol. 100, 5215–5229. doi:10.1007/s00253-016-7520-x

24. Dietrich, L.E., Price-Whelan, A., Petersen, A., Whiteley, M., Newman, D.K., 2006. The phenazine pyocyanin is a terminal signalling factor in the quorum sensing network of *Pseudomonas aeruginosa*. Mol. Microbiol. 61, 1308–1321. doi:10.1111/j.1365-2958.2006.05306.x

25. Dietrich, L.E.P., Okegbe, C., Price-Whelan, A., Sakhtah, H., Hunter, R.C., Newman, D.K., 2013. Bacterial community morphogenesis is intimately linked to the intracellular redox state. J. Bacteriol. 195, 1371–1380. doi:10.1128/JB.02273-12

26. Dietrich, L.E.P., Teal, T.K., Price-Whelan, A., Newman, D.K., 2008. Redox-active antibiotics control gene expression and community behavior in divergent bacteria. Science 321, 1203–1206. doi:10.1126/science.1160619

27. Driscoll, J.A., Brody, S.L., Kollef, M.H., 2007. The epidemiology, pathogenesis and treatment of *Pseudomonas aeruginosa* infections. Drugs 67, 351–368. doi:10.2165/00003495-200767030-00003

28. Du, D., Wang-Kan, X., Neuberger, A., van Veen, H.W., Pos, K.M., Piddock, L.J.V., Luisi, B.F., 2018. Multidrug efflux pumps: structure, function and regulation. Nat. Rev. Microbiol. 16, 523–539. doi:10.1038/s41579-018-0048-6

29. Dubarry, N., Du, W., Lane, D., Pasta, F., 2010. Improved electrotransformation and decreased antibiotic resistance of the cystic fibrosis pathogen *Burkholderia cenocepacia* strain J2315. Appl. Environ. Microbiol. 76, 1095–1102. doi:10.1128/AEM.02123-09

30. Earle, K.A., Billings, G., Sigal, M., Lichtman, J.S., Hansson, G.C., Elias, J.E., Amieva, M.R., Huang, K.C., Sonnenburg, J.L., 2015. Quantitative imaging of gut microbiota spatial organization. Cell Host Microbe 18, 478–488. doi:10.1016/j.chom.2015.09.002

31. EUCAST, E.C. for A.S.T., 2003. Determination of minimum inhibitory concentrations (MICs) of antibacterial agents by broth dilution. Clin. Microbiol. Infect. 9, ix–xv. doi:10.1046/j.1469-0691.2003.00790.x

32. Fair, R.J., Tor, Y., 2014. Antibiotics and bacterial resistance in the 21st century. Perspect. Medicin. Chem. 6, 25–64. doi:10.4137/PMC.S14459

33. Folescu, T.W., da Costa, C.H., Cohen, R.W.F., da Conceição Neto, O.C., Albano, R.M., Marques, E.A., 2015. *Burkholderia cepacia* complex: clinical course in cystic fibrosis patients. BMC Pulm Med 15, 158. doi:10.1186/s12890-015-0148-2

34. Garg, N., Wang, M., Hyde, E., da Silva, R.R., Melnik, A.V., Protsyuk, I., Bouslimani, A., Lim, Y.W., Wong, R., Humphrey, G., Ackermann, G., Spivey, T., Brouha, S.S., Bandeira, N., Lin, G.Y., Rohwer, F., Conrad, D.J., Alexandrov, T., Knight, R., Dorrestein, P.C., 2017. Three-dimensional microbiome and metabolome cartography of a diseased human lung. Cell Host Microbe 22, 705–716.e4. doi:10.1016/j.chom.2017.10.001

35. Gibson, D.G., Young, L., Chuang, R.-Y., Venter, J.C., Hutchison, C.A., Smith, H.O., 2009. Enzymatic assembly of DNA molecules up to several hundred kilobases. Nat. Methods 6, 343–345. doi:10.1038/nmeth.1318

36. Glasser, N.R., Kern, S.E., Newman, D.K., 2014. Phenazine redox cycling enhances anaerobic survival in *Pseudomonas aeruginosa* by facilitating generation of ATP and a proton-motive force. Mol. Microbiol. 92, 399–412. doi:10.1111/mmi.12566

37. Glasser, N.R., Saunders, S.H., Newman, D.K., 2017. The colorful world of extracellular electron shuttles. Annu. Rev. Microbiol. 71, 731–751. doi:10.1146/annurev-micro-090816-093913

38. Graves, M., Robin, T., Chipman, A.M., Wong, J., Khashe, S., Janda, J.M., 1997. Four additional cases of *Burkholderia gladioli* infection with microbiological correlates and review. Clin. Infect. Dis. 25, 838–842. doi:10.1086/515551

39. Gu, M., Imlay, J.A., 2011. The SoxRS response of *Escherichia coli* is directly activated by redox-cycling drugs rather than by superoxide. Mol. Microbiol. 79, 1136–1150. doi:10.1111/j.1365-2958.2010.07520.x

40. Gurevich, A., Saveliev, V., Vyahhi, N., Tesler, G., 2013. QUAST: quality assessment tool for genome assemblies. Bioinformatics 29, 1072–1075. doi:10.1093/bioinformatics/btt086

41. Ham, J.H., Melanson, R.A., Rush, M.C., 2011. *Burkholderia glumae*: next major pathogen of rice? Mol. Plant Pathol. 12, 329–339. doi:10.1111/j.1364-3703.2010.00676.x

42. Harris, C.R., Millman, K.J., van der Walt, S.J., Gommers, R., Virtanen, P., Cournapeau, D., Wieser, E., Taylor, J., Berg, S., Smith, N.J., Kern, R., Picus, M., Hoyer, S., van Kerkwijk, M.H., Brett, M., Haldane, A., Del Río, J.F., Wiebe, M., Peterson, P., Gérard-Marchant, P., Sheppard, K., Reddy, T., Weckesser, W., Abbasi, H., Gohlke, C., Oliphant, T.E., 2020. Array programming with NumPy. Nature 585, 357–362. doi:10.1038/s41586-020-2649-2

43. Hoang, T.T., Karkhoff-Schweizer, R.R., Kutchma, A.J., Schweizer, H.P., 1998. A broad-host-range Flp-FRT recombination system for site-specific excision of chromosomally-located DNA sequences: application for isolation of unmarked *Pseudomonas aeruginosa* mutants. Gene 212, 77–86. doi:10.1016/S0378-1119(98)00130-9

44. Huang, W., Wilks, A., 2017. A rapid seamless method for gene knockout in *Pseudomonas aeruginosa*. BMC Microbiol. 17, 199. doi:10.1186/s12866-017-1112-5

45. Hunter, J.D., 2007. Matplotlib: A 2D Graphics Environment. Comput Sci Eng 9, 90–95. doi:10.1109/MCSE.2007.55

46. Hutchings, M.I., Truman, A.W., Wilkinson, B., 2019. Antibiotics: past, present and future. Curr. Opin. Microbiol. 51, 72–80. doi:10.1016/j.mib.2019.10.008

47. Imlay, J.A., 2013. The molecular mechanisms and physiological consequences of oxidative stress: lessons from a model bacterium. Nat. Rev. Microbiol. 11, 443–454. doi:10.1038/nrmicro3032

48. Jacob, C., Jamier, V., Ba, L.A., 2011. Redox active secondary metabolites. Curr. Opin. Chem. Biol. 15, 149–155. doi:10.1016/j.cbpa.2010.10.015

49. Jeong, Y., Kim, J., Kim, S., Kang, Y., Nagamatsu, T., Hwang, I., 2003. Toxoflavin produced by *Burkholderia glumae* causing rice grain rot is responsible for inducing bacterial wilt in many field crops. Plant Dis. 87, 890–895. doi:10.1094/PDIS.2003.87.8.890

50. Jo, J., Cortez, K.L., Cornell, W.C., Price-Whelan, A., Dietrich, L.E., 2017. An orphan cbb3-type cytochrome oxidase subunit supports *Pseudomonas aeruginosa* biofilm growth and virulence. Elife 6, e30205. doi:10.7554/eLife.30205

51. Jones, C., Webster, G., Mullins, A.J., Jenner, M., Bull, M.J., Dashti, Y., Spilker, T., Parkhill, J., Connor, T.R., LiPuma, J.J., Challis, G.L., Mahenthiralingam, E., 2021. Kill and cure: genomic phylogeny and bioactivity of *Burkholderia gladioli* bacteria capable of pathogenic and beneficial lifestyles. Microb. Genom. 7. doi:10.1099/mgen.0.000515

52. Jorgensen, J.H., Ferraro, M.J., 2009. Antimicrobial susceptibility testing: a review of general principles and contemporary practices. Clin. Infect. Dis. 49, 1749–1755. doi:10.1086/647952

53. Katoh, K., Misawa, K., Kuma, K., Miyata, T., 2002. MAFFT: a novel method for rapid multiple sequence alignment based on fast Fourier transform. Nucleic Acids Res. 30, 3059–3066. doi:10.1093/nar/gkf436

54. Katoh, K., Standley, D.M., 2013. MAFFT multiple sequence alignment software version 7: improvements in performance and usability. Mol. Biol. Evol. 30, 772–780. doi:10.1093/molbev/mst010

55. Kennedy, M.P., Coakley, R.D., Donaldson, S.H., Aris, R.M., Hohneker, K., Wedd, J.P., Knowles, M.R., Gilligan, P.H., Yankaskas, J.R., 2007. *Burkholderia gladioli*: five year experience in a cystic fibrosis and lung transplantation center. J Cyst Fibros 6, 267–273. doi:10.1016/j.jcf.2006.10.007

56. Kester, J.C., Fortune, S.M., 2014. Persisters and beyond: mechanisms of phenotypic drug resistance and drug tolerance in bacteria. Crit Rev Biochem Mol Biol 49, 91–101. doi:10.3109/10409238.2013.869543

57. Kim, J., Kim, J.-G., Kang, Y., Jang, J.Y., Jog, G.J., Lim, J.Y., Kim, S., Suga, H., Nagamatsu, T., Hwang, I., 2004. Quorum sensing and the LysR-type transcriptional activator ToxR regulate toxoflavin biosynthesis and transport in *Burkholderia glumae*. Mol. Microbiol. 54, 921–934. doi:10.1111/j.1365-2958.2004.04338.x

58. Latuasan, H.E., Berends, W., 1961. On the origin of the toxicity of toxoflavin. Biochim. Biophys. Acta 52, 502–508. doi:10.1016/0006-3002(61)90408-5

59. Lau, G.W., Hassett, D.J., Ran, H., Kong, F., 2004. The role of pyocyanin in *Pseudomonas aeruginosa* infection. Trends Mol. Med. 10, 599–606. doi:10.1016/j.molmed.2004.10.002

60. Laursen, J.B., Nielsen, J., 2004. Phenazine natural products: biosynthesis, synthetic analogues, and biological activity. Chem. Rev. 104, 1663–1686. doi:10.1021/cr020473j

61. Lee, J., Park, J., Kim, S., Park, I., Seo, Y.-S., 2016. Differential regulation of toxoflavin production and its role in the enhanced virulence of *Burkholderia gladioli*. Mol. Plant Pathol. 17, 65–76. doi:10.1111/mpp.12262

62. Lee, K.-L., Singh, A.K., Heo, L., Seok, C., Roe, J.-H., 2015. Factors affecting redox potential and differential sensitivity of SoxR to redox-active compounds. Mol. Microbiol. 97, 808– 821. doi:10.1111/mmi.13068

63. Lelis, T., Peng, J., Barphagha, I., Chen, R., Ham, J.H., 2019. The virulence function and regulation of the metalloprotease gene *prtA* in the plant-pathogenic bacterium *Burkholderia glumae*. Mol. Plant Microbe Interact. 32, 841–852. doi:10.1094/MPMI-11-18-0312-R

64. Levin-Reisman, I., Ronin, I., Gefen, O., Braniss, I., Shoresh, N., Balaban, N.Q., 2017. Antibiotic tolerance facilitates the evolution of resistance. Science 355, 826–830. doi:10.1126/science.aaj2191

65. Li, X.-Z., Plésiat, P., Nikaido, H., 2015. The challenge of efflux-mediated antibiotic resistance in Gram-negative bacteria. Clin. Microbiol. Rev. 28, 337–418. doi:10.1128/CMR.00117-14

66. Lipuma, J.J., 2010. The changing microbial epidemiology in cystic fibrosis. Clin. Microbiol. Rev. 23, 299–323. doi:10.1128/CMR.00068-09

67. Liu, G.Y., Nizet, V., 2009. Color me bad: microbial pigments as virulence factors. Trends Microbiol. 17, 406–413. doi:10.1016/j.tim.2009.06.006

68. MacLean, R.C., San Millan, A., 2019. The evolution of antibiotic resistance. Science 365, 1082– 1083. doi:10.1126/science.aax3879

69. Maier, L., Goemans, C.V., Wirbel, J., Kuhn, M., Eberl, C., Pruteanu, M., Müller, P., Garcia-Santamarina, S., Cacace, E., Zhang, B., Gekeler, C., Banerjee, T., Anderson, E.E., Milanese, A., Löber, U., Forslund, S.K., Patil, K.R., Zimmermann, M., Stecher, B., Zeller, G., Bork, P., Typas, A., 2021. Unravelling the collateral damage of antibiotics on gut bacteria. Nature 599, 120–124. doi:10.1038/s41586-021-03986-2

70. Maier, L., Pruteanu, M., Kuhn, M., Zeller, G., Telzerow, A., Anderson, E.E., Brochado, A.R., Fernandez, K.C., Dose, H., Mori, H., Patil, K.R., Bork, P., Typas, A., 2018. Extensive impact of non-antibiotic drugs on human gut bacteria. Nature 555, 623–628. doi:10.1038/nature25979

71. Martinez, J.L., 2009. The role of natural environments in the evolution of resistance traits in pathogenic bacteria. Proc. Biol. Sci. 276, 2521–2530. doi:10.1098/rspb.2009.0320

72. McKinney, W., 2010. Data structures for statistical computing in python, in: Proceedings of the 9th Python in Science Conference, Proceedings of the Python in Science Conference. Presented at the Python in Science Conference, SciPy, pp. 56–61. doi:10.25080/Majora-92bf1922-00a

73. Meirelles, L.A., Newman, D.K., 2018. Both toxic and beneficial effects of pyocyanin contribute to the lifecycle of *Pseudomonas aeruginosa*. Mol. Microbiol. 110, 995–1010. doi:10.1111/mmi.14132

74. Meirelles, L.A., Perry, E.K., Bergkessel, M., Newman, D.K., 2021. Bacterial defenses against a natural antibiotic promote collateral resilience to clinical antibiotics. PLoS Biol. 19, e3001093. doi:10.1371/journal.pbio.3001093

75. Mukherjee, S., Stamatis, D., Bertsch, J., Ovchinnikova, G., Sundaramurthi, J.C., Lee, J., Kandimalla, M., Chen, I.-M.A., Kyrpides, N.C., Reddy, T.B.K., 2021. Genomes OnLine Database (GOLD) v.8: overview and updates. Nucleic Acids Res. 49, D723–D733. doi:10.1093/nar/gkaa983

76. O’Neill, J., 2016. Tackling drug-resistant infections globally: final report and recommendations.

77. Perry, E.K., Meirelles, L.A., Newman, D.K., 2021. From the soil to the clinic: the impact of microbial secondary metabolites on antibiotic tolerance and resistance. Nat. Rev. Microbiol. doi:10.1038/s41579-021-00620-w

78. Podnecky, N.L., Rhodes, K.A., Schweizer, H.P., 2015. Efflux pump-mediated drug resistance in *Burkholderia*. Front. Microbiol. 6, 305. doi:10.3389/fmicb.2015.00305

79. Powell, D.R., 2015. Degust: interactive RNA-seq analysis. Zenodo. doi:10.5281/zenodo.3258932

80. Price-Whelan, A., Dietrich, L.E.P., Newman, D.K., 2006. Rethinking “secondary” metabolism: physiological roles for phenazine antibiotics. Nat. Chem. Biol. 2, 71–78. doi:10.1038/nchembio764

81. Radlinski, L., Rowe, S.E., Kartchner, L.B., Maile, R., Cairns, B.A., Vitko, N.P., Gode, C.J., Lachiewicz, A.M., Wolfgang, M.C., Conlon, B.P., 2017. *Pseudomonas aeruginosa* exoproducts determine antibiotic efficacy against *Staphylococcus aureus*. PLoS Biol. 15, e2003981. doi:10.1371/journal.pbio.2003981

82. Ramos, I., Dietrich, L.E., Price-Whelan, A., Newman, D.K., 2010. Phenazines affect biofilm formation by *Pseudomonas aeruginosa* in similar ways at various scales. Res. Microbiol. 161, 187–191. doi:10.1016/j.resmic.2010.01.003

83. Reszka, K.J., O’Malley, Y., McCormick, M.L., Denning, G.M., Britigan, B.E., 2004. Oxidation of pyocyanin, a cytotoxic product from *Pseudomonas aeruginosa*, by microperoxidase 11 and hydrogen peroxide. Free Radic. Biol. Med. 36, 1448–1459. doi:10.1016/j.freeradbiomed.2004.03.011

84. Rhodes, K.A., Schweizer, H.P., 2016. Antibiotic resistance in *Burkholderia* species. Drug Resist Updat 28, 82–90. doi:10.1016/j.drup.2016.07.003

85. Ronquist, F., Teslenko, M., van der Mark, P., Ayres, D.L., Darling, A., Höhna, S., Larget, B., Liu, L., Suchard, M.A., Huelsenbeck, J.P., 2012. MrBayes 3.2: efficient Bayesian phylogenetic inference and model choice across a large model space. Syst. Biol. 61, 539–542. doi:10.1093/sysbio/sys029

86. Saunders, S.H., Tse, E.C.M., Yates, M.D., Otero, F.J., Trammell, S.A., Stemp, E.D.A., Barton, J.K., Tender, L.M., Newman, D.K., 2020. Extracellular DNA promotes efficient extracellular electron transfer by pyocyanin in *Pseudomonas aeruginosa* biofilms. Cell 182, 919–932.e19. doi:10.1016/j.cell.2020.07.006

87. Schiessl, K.T., Hu, F., Jo, J., Nazia, S.Z., Wang, B., Price-Whelan, A., Min, W., Dietrich, L.E.P., 2019. Phenazine production promotes antibiotic tolerance and metabolic heterogeneity in *Pseudomonas aeruginosa* biofilms. Nat. Commun. 10, 762. doi:10.1038/s41467-019-08733-w

88. Schnetterle M, Gorgé O, Nolent F, Boughammoura A, Sarilar V, Vigier C, Guillier S, Koch L, Degand N, Ramisse V, Tichadou X, Girleanu M, Favier A-L, Valade E, Biot F, Neulat-Ripoll F. 2021. Genomic and RT-qPCR analysis of trimethoprim-sulfamethoxazole and meropenem resistance in *Burkholderia pseudomallei* clinical isolates. PLoS Negl Trop Dis 15:e0008913. doi:10.1371/journal.pntd.0008913

89. Schwab, U., Abdullah, L.H., Perlmutt, O.S., Albert, D., Davis, C.W., Arnold, R.R., Yankaskas, J.R., Gilligan, P., Neubauer, H., Randell, S.H., Boucher, R.C., 2014. Localization of *Burkholderia cepacia* complex bacteria in cystic fibrosis lungs and interactions with *Pseudomonas aeruginosa* in hypoxic mucus. Infect. Immun. 82, 4729–4745. doi:10.1128/IAI.01876-14

90. Seemann, T., 2014. Prokka: rapid prokaryotic genome annotation. Bioinformatics 30, 2068– 2069. doi:10.1093/bioinformatics/btu153

91. Sheplock, R., Recinos, D.A., Mackow, N., Dietrich, L.E.P., Chander, M., 2013. Species-specific residues calibrate SoxR sensitivity to redox-active molecules. Mol. Microbiol. 87, 368– 381. doi:10.1111/mmi.12101

92. Shi, H., Shi, Q., Grodner, B., Lenz, J.S., Zipfel, W.R., Brito, I.L., De Vlaminck, I., 2020. Highly multiplexed spatial mapping of microbial communities. Nature 588, 676–681. doi:10.1038/s41586-020-2983-4

93. Simon, R., Priefer, U., Pühler, A., 1983. A broad host range mobilization system for *in vivo* genetic engineering: transposon mutagenesis in gram negative bacteria. Nat. Biotechnol. 1, 784–791. doi:10.1038/nbt1183-784

94. Singh, A.K., Shin, J.-H., Lee, K.-L., Imlay, J.A., Roe, J.-H., 2013. Comparative study of SoxR activation by redox-active compounds. Mol. Microbiol. 90, 983–996. doi:10.1111/mmi.12410

95. Skinnider, M.A., Johnston, C.W., Gunabalasingam, M., Merwin, N.J., Kieliszek, A.M., MacLellan, R.J., Li, H., Ranieri, M.R.M., Webster, A.L.H., Cao, M.P.T., Pfeifle, A., Spencer, N., To, Q.H., Wallace, D.P., Dejong, C.A., Magarvey, N.A., 2020. Comprehensive prediction of secondary metabolite structure and biological activity from microbial genome sequences. Nat. Commun. 11, 6058. doi:10.1038/s41467-020-19986-1

96. Stern, K.G., 1935. Oxidation-reduction potentials of toxoflavin. Biochem. J. 29, 500–508. doi:10.1042/bj0290500

97. The pandas development team, 2020. pandas-dev/pandas: Pandas 1.0.3. Zenodo. doi:10.5281/zenodo.3509134

98. Tjaden, B., 2015. De novo assembly of bacterial transcriptomes from RNA-seq data. Genome Biol. 16, 1. doi:10.1186/s13059-014-0572-2

99. Tyc, O., Song, C., Dickschat, J.S., Vos, M., Garbeva, P., 2017. The ecological role of volatile and soluble secondary metabolites produced by soil bacteria. Trends Microbiol. 25, 280– 292. doi:10.1016/j.tim.2016.12.002

100. Vernocchi, P., Del Chierico, F., Putignani, L., 2016. Gut microbiota profiling: metabolomics based approach to unravel compounds affecting human health. Front. Microbiol. 7, 1144. doi:10.3389/fmicb.2016.01144

101. Waglechner, N., McArthur, A.G., Wright, G.D., 2019. Phylogenetic reconciliation reveals the natural history of glycopeptide antibiotic biosynthesis and resistance. Nat. Microbiol. 4, 1862–1871. doi:10.1038/s41564-019-0531-5

102. Waskom, M., 2021. seaborn: statistical data visualization. JOSS 6, 3021. doi:10.21105/joss.03021

103. Wick, R.R., Judd, L.M., Gorrie, C.L., Holt, K.E., 2017. Unicycler: Resolving bacterial genome assemblies from short and long sequencing reads. PLoS Comput. Biol. 13, e1005595. doi:10.1371/journal.pcbi.1005595

104. Widdel, F., Pfennig, N., 1981. Studies on dissimilatory sulfate-reducing bacteria that decompose fatty acids. Arch. Microbiol. 129, 395–400. doi:10.1007/BF00406470

105. Wilbert, S.A., Mark Welch, J.L., Borisy, G.G., 2020. Spatial ecology of the human tongue dorsum microbiome. Cell Rep. 30, 4003–4015.e3. doi:10.1016/j.celrep.2020.02.097

106. Wilson, R., Sykes, D.A., Watson, D., Rutman, A., Taylor, G.W., Cole, P.J., 1988. Measurement of *Pseudomonas aeruginosa* phenazine pigments in sputum and assessment of their contribution to sputum sol toxicity for respiratory epithelium. Infect. Immun. 56, 2515– 2517. doi:10.1128/IAI.56.9.2515-2517.1988

107. Windels, E.M., Michiels, J.E., Fauvart, M., Wenseleers, T., Van den Bergh, B., Michiels, J., 2019a. Bacterial persistence promotes the evolution of antibiotic resistance by increasing survival and mutation rates. ISME J. 13, 1239–1251. doi:10.1038/s41396-019-0344-9

108. Windels, E.M., Michiels, J.E., Van den Bergh, B., Fauvart, M., Michiels, J., 2019b. Antibiotics: combatting tolerance to stop resistance. MBio 10. doi:10.1128/mBio.02095-19

109. Winsor, G.L., Griffiths, E.J., Lo, R., Dhillon, B.K., Shay, J.A., Brinkman, F.S.L., 2016. Enhanced annotations and features for comparing thousands of *Pseudomonas* genomes in the *Pseudomonas* genome database. Nucleic Acids Res. 44, D646–53. doi:10.1093/nar/gkv1227

110. Winsor, G.L., Khaira, B., Van Rossum, T., Lo, R., Whiteside, M.D., Brinkman, F.S.L., 2008. The *Burkholderia* Genome Database: facilitating flexible queries and comparative analyses. Bioinformatics 24, 2803–2804. doi:10.1093/bioinformatics/btn524

111. Zhu, K., Chen, S., Sysoeva, T.A., You, L., 2019. Universal antibiotic tolerance arising from antibiotic-triggered accumulation of pyocyanin in *Pseudomonas aeruginosa*. PLoS Biol. 17, e3000573. doi:10.1371/journal.pbio.3000573

112. Zimmermann, M., Patil, K.R., Typas, A., Maier, L., 2021. Towards a mechanistic understanding of reciprocal drug-microbiome interactions. Mol. Syst. Biol. 17, e10116. doi:10.15252/msb.202010116

