## Supplementary figures and Table legends for "Redox-active secondary metabolites act as interspecies modulators of antibiotic resilience"

### SUPPLEMENTARY INFORMATION

#### Supplementary Figures

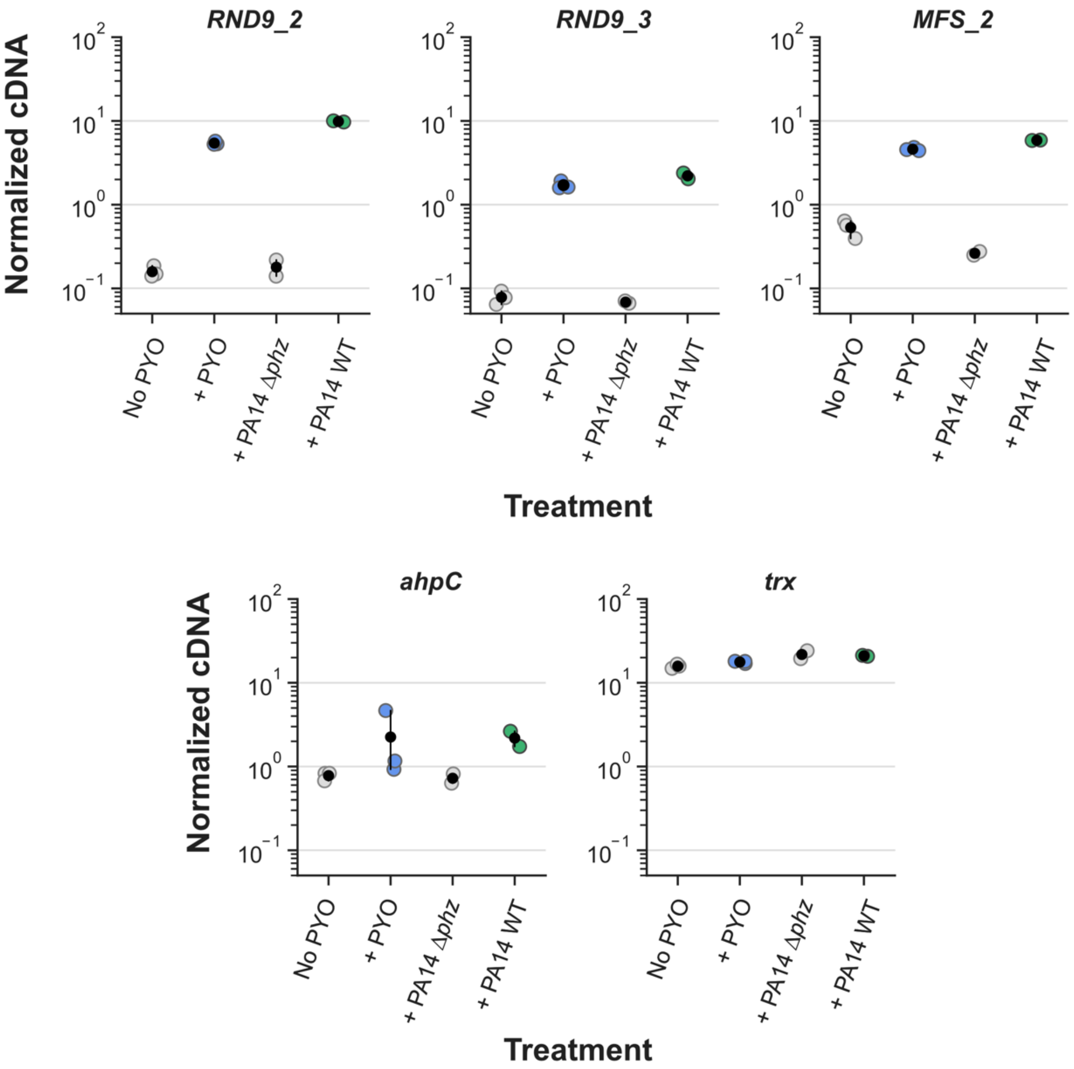


**Figure S1. Results for qRT-PCR experiment 1 (normalized cDNA levels).** Data for each gene show in Fig. 1C-D, displayed as normalized cDNA for comparison. Normalizations were done by the housekeeping gene *uvrC* (see Materials and Methods). Data for genes RND9_2 and RND9_3 (only for *B. multivorans* treatments with and without PYO) is also shown in Fig. 2B and Fig. S2, respectively, for comparison with other *B. multivorans* strains. Black dots mark the means and error bars represent 95% confidence intervals.

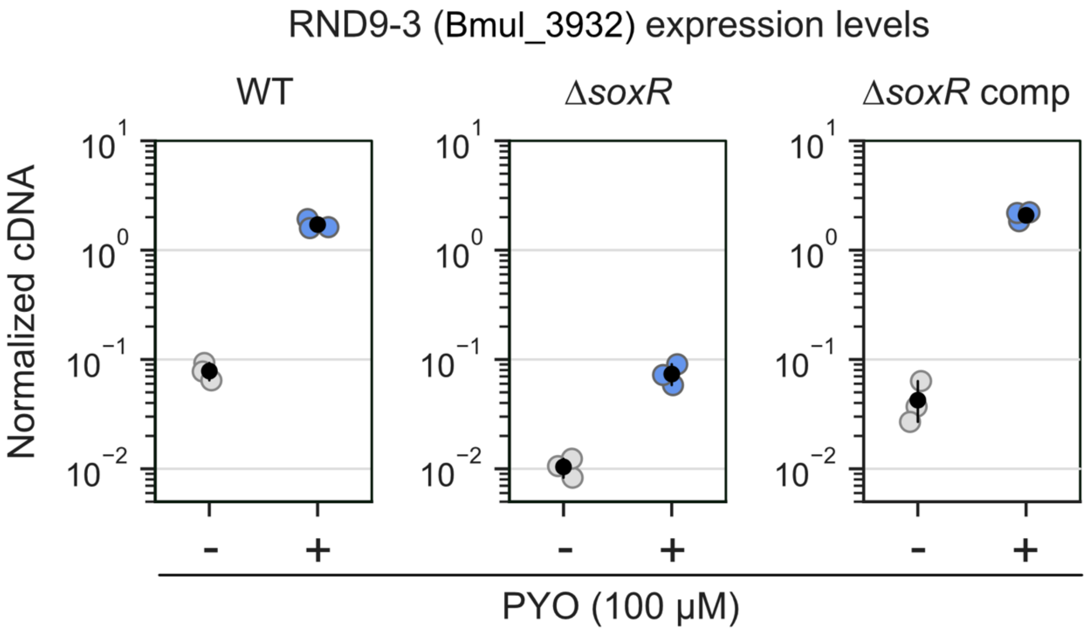


**Figure S2. Expression levels of the third gene in the RND-9 operon (Bmul_3932) in the presence or absence of PYO**. Values measured by qRT-PCR in different *B. multivorans* strains (n = 3). Δ*soxR* comp means complementation of Δ*soxR*. Data is shown as normalized cDNA. Normalizations were done by the housekeeping gene *uvrC* (see Materials and Methods). Data for the *B. multivoran*s WT strain is the same as in the experiment displayed in Fig. S1 but is shown here for comparison. Black dots mark the means and error bars represent 95% confidence intervals.

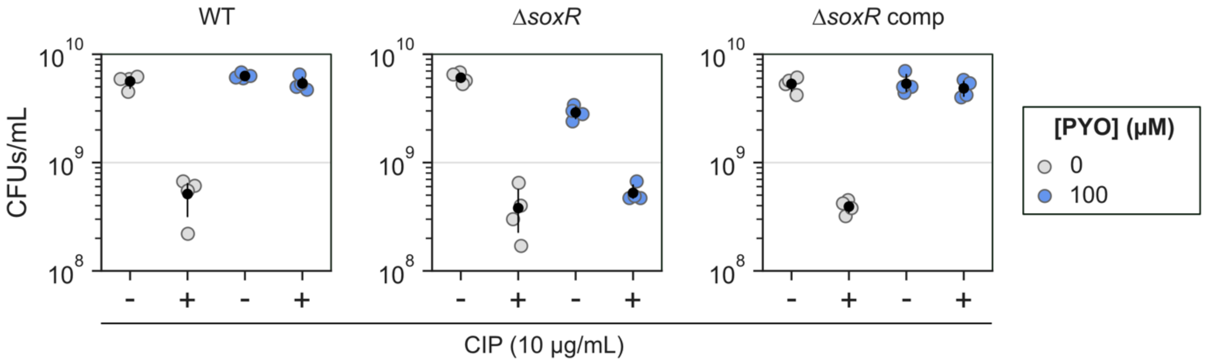


**Figure S3. Raw colony forming unit (CFU) numbers during ciprofloxacin (CIP) tolerance experiments with WT, Δ*soxR* and Δ*soxR* comp strains**. In these experiments, the No CIP controls ("-") were used as the baseline for calculation of survival rates displayed in Fig. 2C (n = 4). Notice that the Δ*soxR* strain was more sensitive to PYO than the other strains, visible by its lower CFUs in the "No CIP control + PYO". This resulted in the apparent PYO-mediated slight increase in tolerance for this strain shown for this strain in Fig. 2C. However, comparisons of the raw CFUs after treatment with CIP ("+") show that PYO did not have a meaningful effect on the CIP tolerance of this strain. Black dots mark the means and error bars represent 95% confidence intervals.

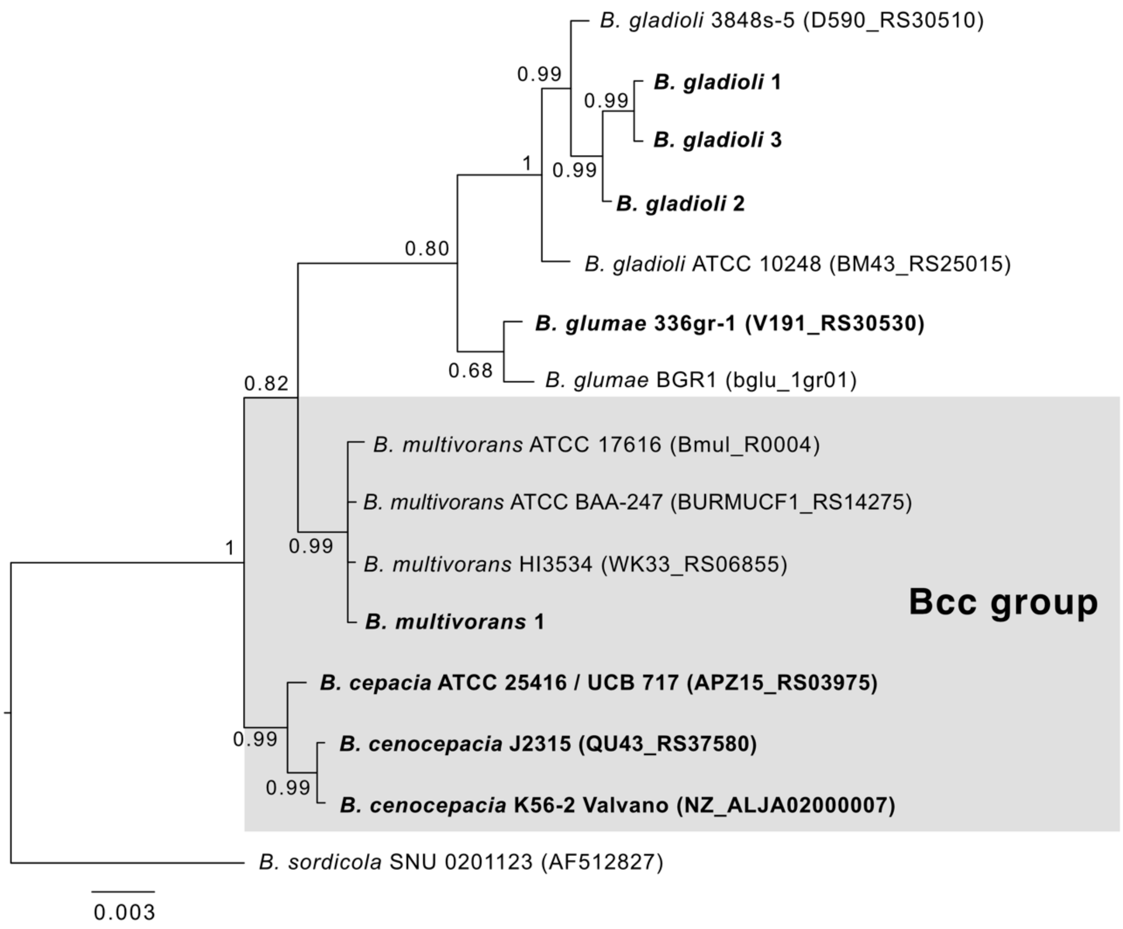


**Figure S4. Full phylogenetic tree of *Burkholderia* species used in the study.** Phylogeny is based on 16S rRNA sequences. Strains highlighted in bold were used in experiments throughout the study, and gray shading highlights species within the Bcc group. Posterior probabilities of Bayesian analysis are shown in the nodes. For sequences retrieved from the BGD database, the respective locus tag is given in parenthesis. *B. sordicula*, a *Burkholderia* species from the "*B. gathei* group" (Depoorter et al., 2016), was used as an outgroup (for this sequence, the number in parenthesis corresponds to its GenBank accession number). The alignment used during the phylogenetic analysis is available as supplementary material.

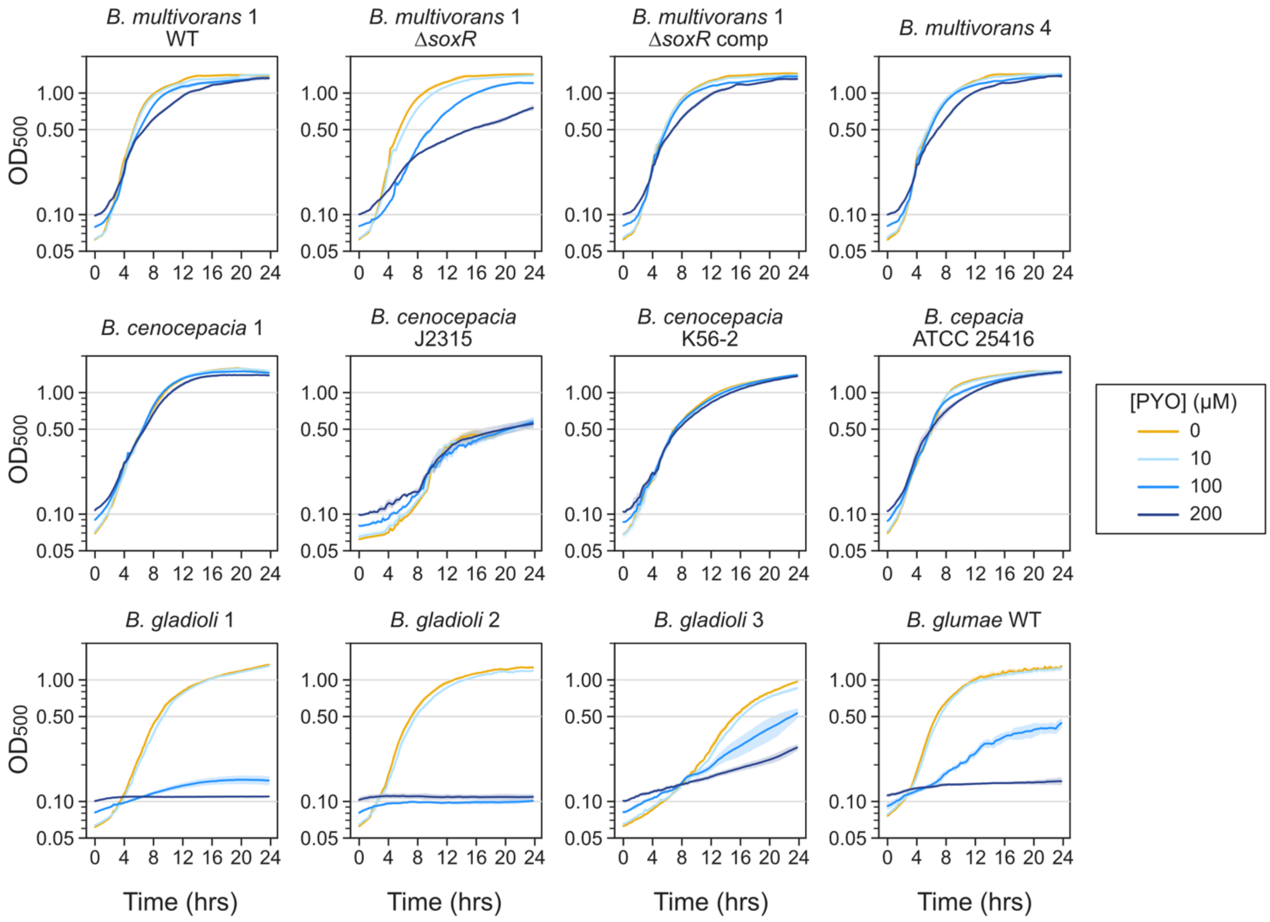


**Figure S5. Growth curves for different *Burkholderia* species (all strains studied are included) under different concentrations of PYO.** Notice accentuated growth defects for *B. gladioli*, *B. glumae* and *B. multivorans* 1 Δ*soxR* under high concentrations of PYO (100 µM or higher). Plotted lines represent means of three replicates, and shaded areas represent the standard deviations. This dataset was used to calculate the growth rates used in Fig. 2E.

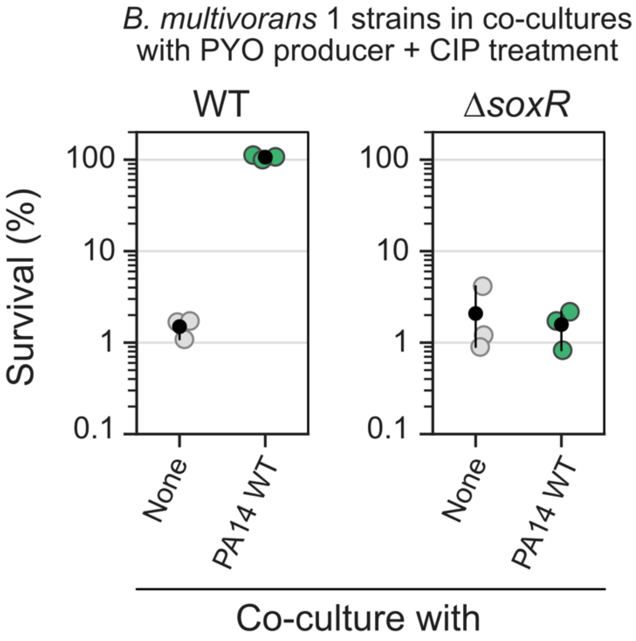


**Figure S6. Tolerance of *B. multivorans* WT and Δ*soxR* when alone or in co-cultures with *P. aeruginosa*.** PA14 = *P. aeruginosa* PA14 (PYO producer). PYO produced by *P. aeruginosa* induced tolerance to CIP (10 µg/mL) in *B. multivorans* 1 WT, but not in *B. multivorans* 1 Δ*soxR* (n = 3). Black dots mark the means and error bars represent 95% confidence intervals.

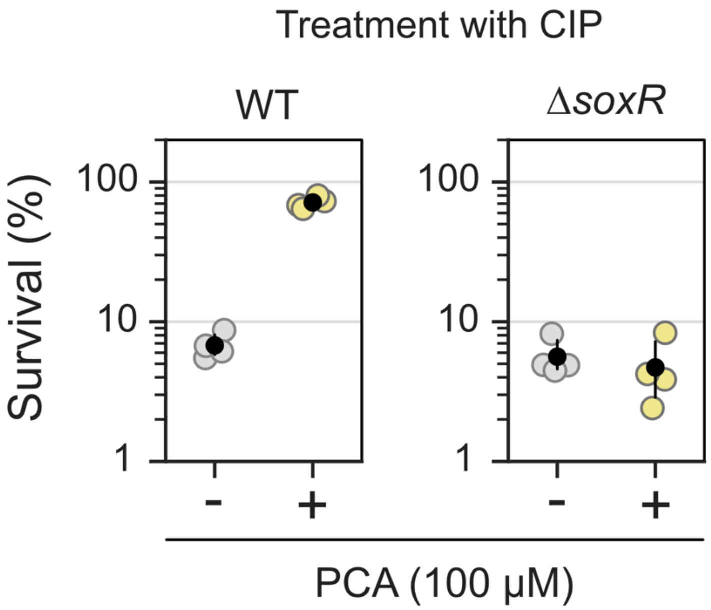


**Figure S7. Effect of phenazine-1-carboxylic acid (PCA) on tolerance in *B. multivorans* 1.** Results for WT and Δ*soxR* strains are shown (n = 4) after exposure to ciprofloxacin (CIP, 10 µg/mL). Black dots mark the means and error bars represent 95% confidence intervals.

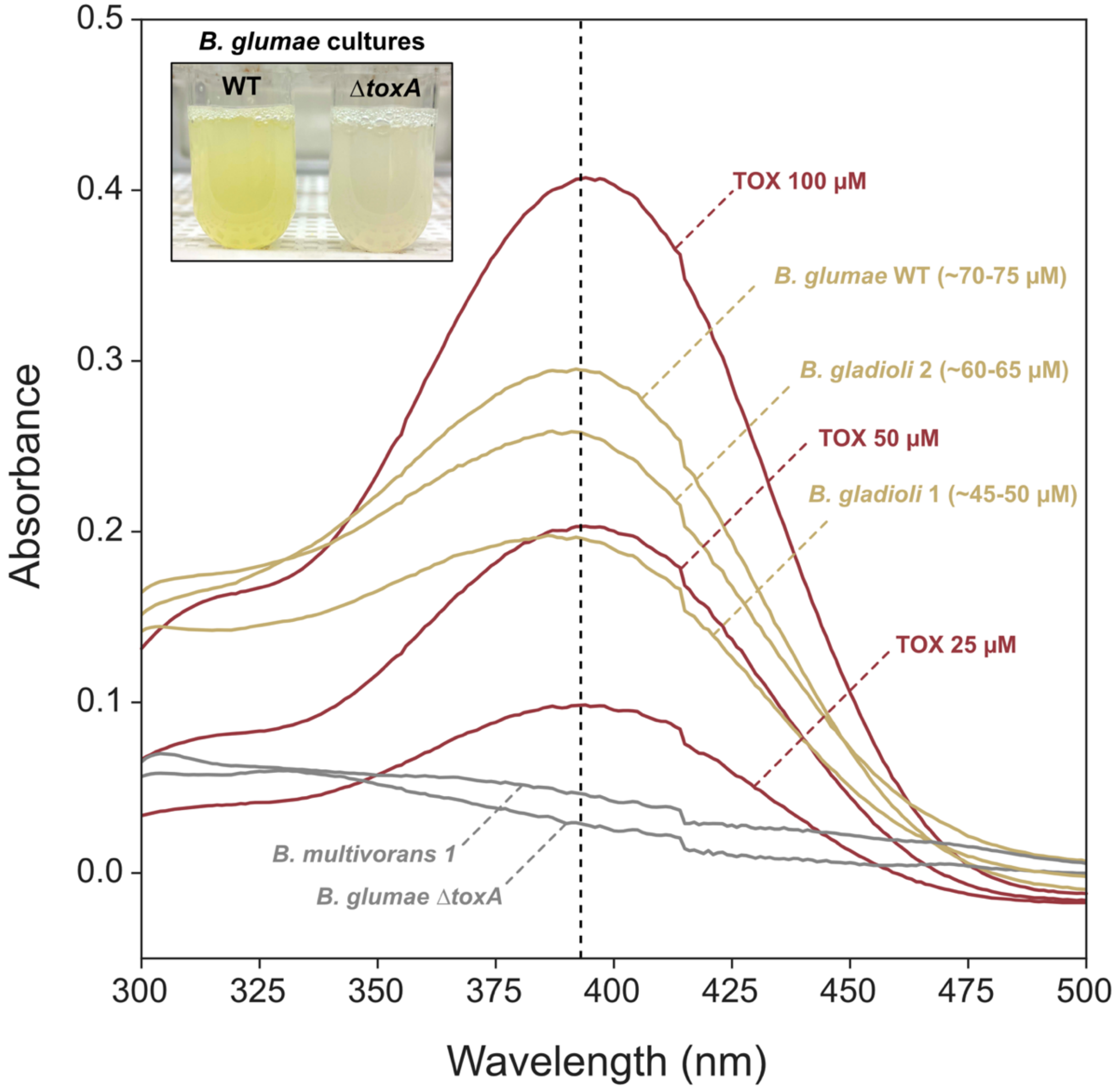


**Figure S8. Toxoflavin (TOX) production by the *B. glumae* and *B. gladioli* species studied.** Absorbance spectra for wavelengths between 300-500 nm are shown for culture supernatants of *Burkholderia* strains together with different standard concentrations of pure TOX. Absorbance at 393 nm (dashed line) is commonly used for TOX quantification (Chen et al., 2012; Kim et al., 2004). Concentrations for cultures are estimations based on standards using the pure molecule. *B. glumae* WT is a validated TOX producer, and the *B. glumae* Δ*toxA* strain cannot make the metabolite (Lelis et al., 2019). Tube pictures on the top left show yellow pigmentation produced by the *B. glumae* WT strain culture in comparison to the Δ*toxA*. Cultures shown in the picture were not the same used during UV-Vis spectra collection. Picture saturation was increased for better visualization of the yellow pigmentation in the tubes.

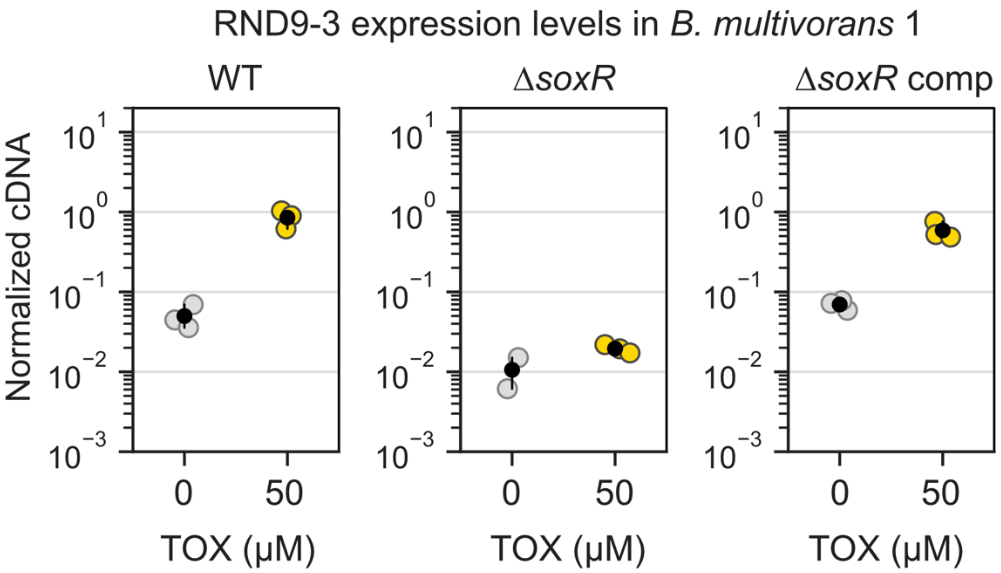


**Figure S9. Expression levels of the third gene in the RND-9 operon (Bmul_3932) in the presence or absence of TOX**. Values measured by qRT-PCR in different *B. multivorans* strains (n = 3 for all except "Δ*soxR* + 0 μM TOX", where n = 2). Δ*soxR* comp means complementation of Δ*soxR*. Data is shown as normalized cDNA. Normalizations were done by the housekeeping gene *uvrC* (see Materials and Methods). Black dots mark the means and error bars represent 95% confidence intervals.

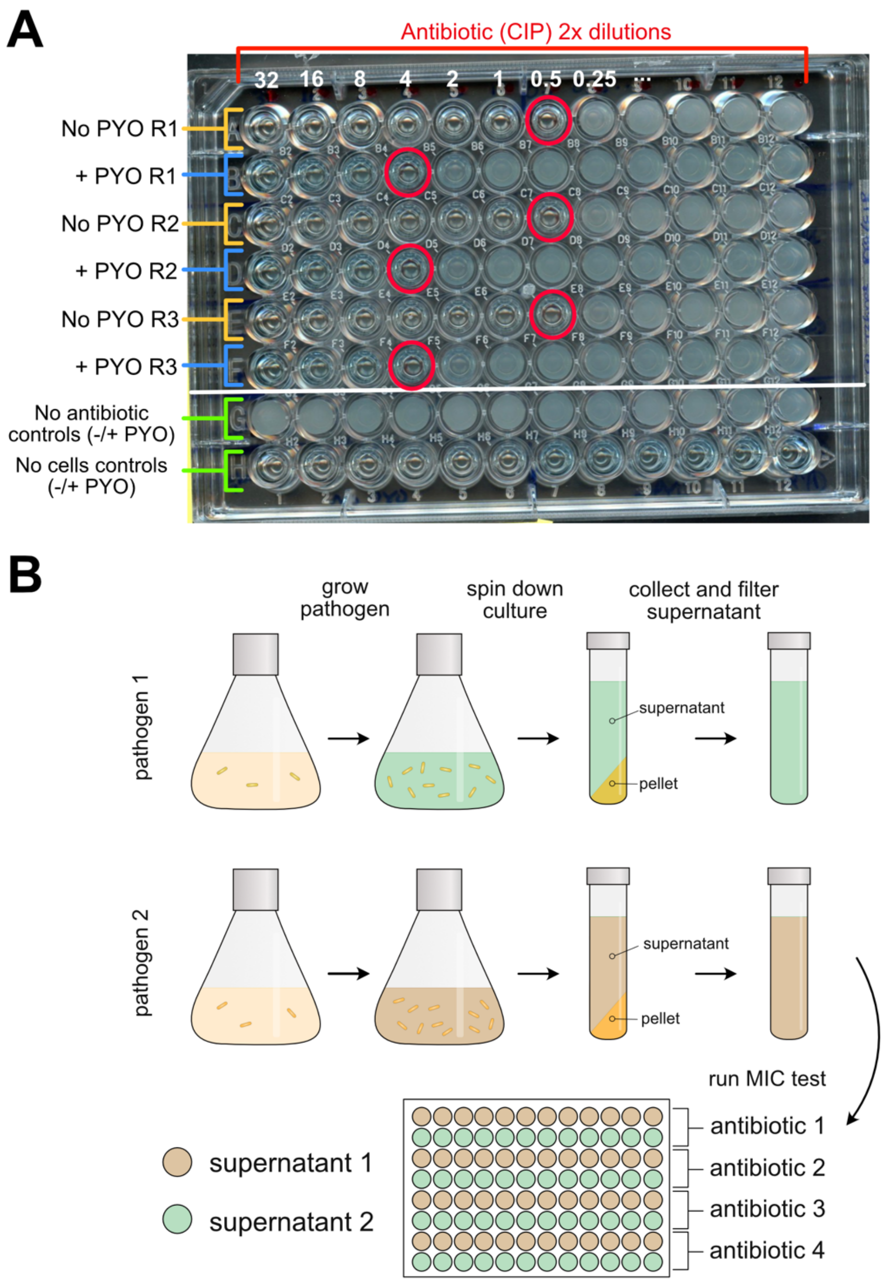


**Figure S10. Additional information on experimental design during MIC assays. A.** Example of experimental plate testing the effect of PYO on *B. multivorans* 1 WT resistance to MIC ciprofloxacin (CIP). Three replicates were prepared (R1-R3), with and without PYO added to the fresh medium. Cells always grew in the "no antibiotic" controls, independently of the presence of PYO (or other secondary metabolite tested). No growth was observed in the "no cells" controls. Red circles marks MICs detected based on analysis as described in the Materials and Methods. **B.** Cartoon describing the protocol for MIC experiments using spent media. Spent media were four-fold diluted into fresh media before the experiments (see Materials and Methods).

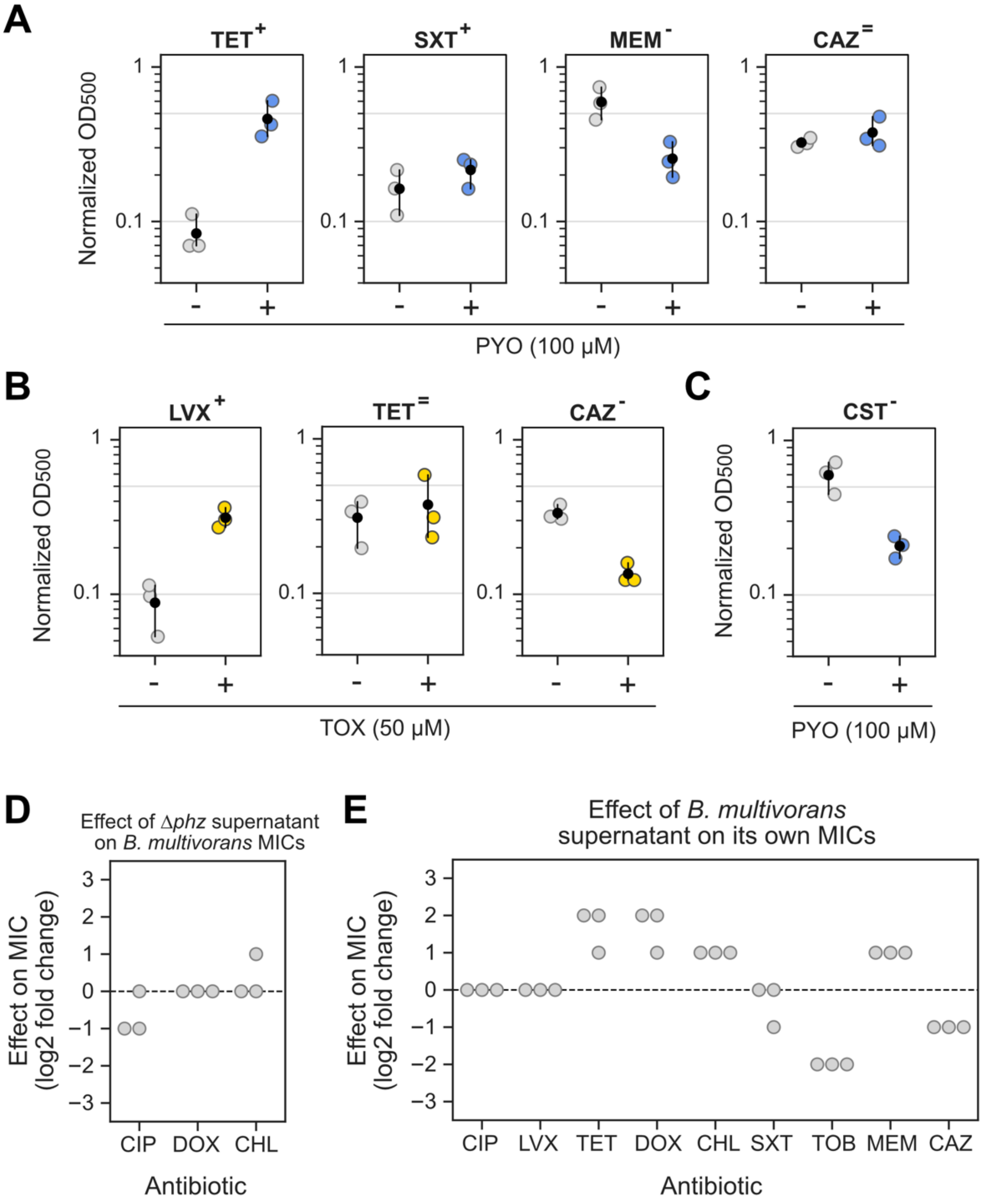


**Figure S11. Additional MIC results. A-B.** Normalized absorbance measurements ($OD_{500}$) used for the calculation of the fold changes on growth density caused by addition of PYO (A) or TOX (B) displayed in Figs. 4D. **C.** Effects of PYO on growth in the presence of colistin (CST, 4,096 µg/mL, a concentration still not enough to completely inhibit growth under these conditions). **D.** Effects of *P. aeruginosa* Δ*phz* supernatant (i.e. no PYO present) on *B. multivorans* 1 MICs for three antibiotics where PYO had increased MICs (for each antibiotic, n = 3). **E.** Effects of *B. multivorans* 1 supernatant on its own MICs (for each antibiotic, n = 3). In panels A-C, the black dots mark the means and error bars represent 95% confidence intervals.

CIP, ciprofloxacin; LVX, levofloxacin; TET, tetracycline, DOX, doxycycline, CHL, chloramphenicol, SXT, sulfamethoxazole/trimethoprim; TOB, tobramycin; MEM, meropenem; CAZ, ceftazidime; CST, colistin.

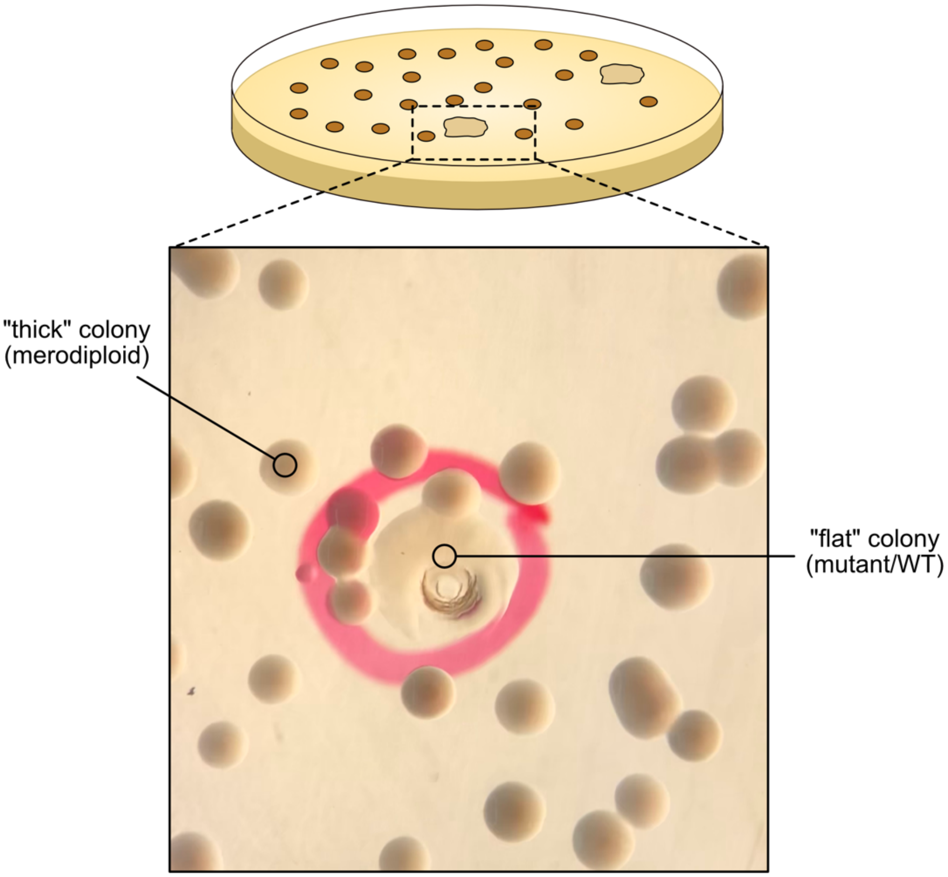


**Figure S12. Colony screen in *B. multivorans* 1 during strains construction.** The desired “flat” colonies were rare (the picture in the bottom part shows one that had been picked for later analysis). These “flat” colonies had lost the construct originally inserted during homologous recombination and had either the mutant or WT genotype, screened by PCR. Abundant “thick” colonies were merodiploids still containing the construct with tetracycline resistance marker integrated in the genome. While we do not know why merodiploids present as round but “cured” strains present as flat, this phenotype is convenient for mutant identification. This screen was done using LB plates (no NaCl) containing only 1.05% agar (see Material and Methods).

**Supplementary Tables Legends**

**Table S1.** Strains, plasmids and primers used in this study.

**Table S2.** Full results of the RNA-seq experiment. See information within each tab for distinct comparisons between treatments. It is also included a tab with the information about the loci tags and gene names used in Fig. 1.

**Table S3.** BLASTP results for the protein sequence of Bmul_3929 (SoxR) from *B. multivorans* 1 against *Pseudomonas aeruginosa* PA14 and PAO1 database of protein sequences.

**Table S4.** MIC values measured in *B. multivorans* 1 for all the antibiotics and conditions tested in the study.
